# Development of a point-of-care field diagnostic test for DFT1 and DFT2

**DOI:** 10.64898/2025.12.17.695037

**Authors:** Anuk Kruawan, K.C. Rajendra, David A Gell, Jocelyn M. Darby, Weizhen Zhu, Chrissie E. B. Ong, Kirsten A. Fairfax, Andrew S. Flies

## Abstract

The Tasmanian devil (*Sarcophilus harrisii*) population has undergone a major decline in the wild due to the epidemics of two transmissible cancers known as devil facial tumours 1 (DFT1) and devil facial tumour 2 (DFT2). Understanding the distribution and prevalence of DFT1 and DFT2 is challenging as they can only be detected in field conditions when tumours are large enough to be visually observed. This impedes our ability to understand how the devil population is evolving in response to the tumours. PCR-based diagnostics are available, but the technique is not field-applicable. To overcome these hurdles, we developed an ultrasensitive and field-deployable rapid diagnostic tool. We used a technique called Specific High-Sensitivity Enzymatic Reporter Unlocking (SHERLOCK), which combines an isothermal nucleic acid amplification step with CRISPR-based recognition and activation of reporter probes to indicate target sequence detection. This system requires only portable laboratory equipment, making it more amenable to field conditions. The results showed that our test can detect DFT1 and DFT2 DNA within the attomolar range. We then confirmed that the test can detect DFT1 in the field using non-invasive swab samples. The SHERLOCK test can be used for more active monitoring and management approaches that are not currently possible.

## INTRODUCTION

The Tasmanian devil (*Sarcophilus harrisii*) is the largest extant carnivorous marsupial and is found exclusively in the island state of Tasmania, Australia [1]. Devils became extinct on mainland Australia around 3000 years ago, possibly due to competition with dingoes (*Canis lupus dingo*), climate change, and human activity [2, 3]. Devils inhabit various regions across Tasmania, including woodlands, eucalyptus forests, and coastal areas [4]. Adult male devils typically weigh between 7.5 and 13 kg, whilst females range from 4.5 to 9 kg [1, 5]. Both sexes can live at least six years in the wild [1, 5]. Tasmanian devils are nocturnal and solitary scavengers, with their home ranges between 12 and 18 km^2^ [6]. Unfortunately, the wild devil population has greatly declined due to two contagious cancers known as devil facial tumour 1 (DFT1) and devil facial tumour 2 (DFT2) [7, 8].

DFT1 and DFT2 appear grossly similar but they are genetically, histologically and cytogenetically different [8, 9]. Various diagnostic assays are available to identify and distinguish the two tumour types, with ‘Tasman PCR’ being the gold standard [10]. Whilst the Tasman PCR can effectively identify DFT1 and DFT2 DNA in tumour biopsies, the technique is challenging to implement in the field. This can lead to diagnosis being made days or weeks after sample collection, limiting further investigation of suspected tumours when devils are in hand during trapping trips. Identification of devils that are exposed to DFT1 or DFT2 that do not develop tumours, develop small tumours but not severe disease, or exhibit natural tumour regressions is critical information for devil conservation management and understanding tumour and devil co-evolution [11–13].

Isothermal amplification assays, such as loop-mediated isothermal amplification (LAMP) and recombinase polymerase amplification (RPA) have been explored as alternative diagnostic methods for a variety of pathogens [14]. LAMP uses eight primers to efficiently amplify the target whilst RPA uses two primers similar to PCR and can operate with PCR primers in some instances [15, 16]. As LAMP requires more primers than RPA, the assay design and optimisation is more challenging [17]. Studies have compared the amplification kinetics among various methods and found that RPA is more rapid and sensitive than LAMP or even PCR [18–20]. However, RPA usually suffers from non-specific amplification and primer dimerisation due to the lower amplification temperature of 37 °C [20]. To improve the specificity of target detection, isothermal amplification techniques can be paired with Clustered Regularly Interspaced Palindromic Repeats (CRISPR) and CRISPR-Cas enzyme for secondary target sequence detection [14].

CRISPR evolved as a defence mechanism in prokaryotes to recognise and degrade pathogen’s (i.e. bacteriophage) genetic materials [21, 22]. The programmable endonuclease activities have been harnessed for gene editing, namely the CRISPR-Cas9 system [23, 24]. Furthermore, other types of Cas enzymes have been used as DNA and RNA detection tools. DNA endonuclease-targeted CRISPR trans reporter (DETECTR) [25] and Specific High-Sensitivity Enzymatic Reporter UnLOCKing (SHERLOCK) [26] are key examples that utilise LbCas12a and LwCas13a enzymes with their corresponding programmable CRISPR RNA (crRNA) to detect DNA and RNA targets, respectively. These techniques incorporate RPA to pre-amplify the target before detection by CRISPR-Cas, which can achieve a single copy sensitivity with high specificity. DETECTR and SHERLOCK have been used to detect pathogens such as SARS-COV-2, *Plasmodium spp*., and food borne diseases such as *Vibrio harveyias* [18, 27, 28]. Moreover, SHERLOCK was the first Food and Drug Administration (FDA)-approved CRISPR-based diagnostic tool used to detect SARS-COV-2 [29].

LbCas12a and LwCas13a recognise single-stranded (ss)DNA or ssRNA, respectively, and both enzymes exhibit collateral cleaving activity (i.e. cleavage of all DNA or RNA in the reaction) upon recognition of the target nucleic acid [25, 26]. SsDNA and ssRNA reporters containing a quencher and fluorophore have been used to detect the collateral cleaving upon target recognition in LbCas12a and LwCas13a, respectively [25, 26]. As LwCas13a only recognises RNA targets, the pre-amplified DNA needs to be converted to RNA, usually through incorporation of a T7 promoter in the forward RPA primer and a T7 polymerase enzyme in the reaction mix, before the CRISPR-Cas cleaving can be activated [30]. In case of RPA-CRISPR-Cas (SHERLOCK) diagnostics, all components only require a single incubation temperature, making it more amenable to field applications than conventional PCR or LAMP-CRISPR (LAMP operates at a higher temperature than CRISPR-Cas) [14].

Here, we describe the development of a SHERLOCK test requiring minimal instrumentation for field diagnosis of DFT1 and DFT2. Our SHERLOCK tests detect interchromosomal fusions identified in DFT1 and DFT2 [10]. The results showed that our test can detect DFT1 and DFT2 DNA within the attomolar DNA range. Pilot field testing revealed that our test could detect DFT1 with a high level of sensitivity and specificity and may have the potential to serve as a preclinical diagnostics for devil facial tumour disease (DFTD).

## MATERIALS AND METHODS

### Cell lines and primary samples

DFT1 C5065 (RRID:CVCL_LB79) [31], DFT2 JV (RRID:CVCL_A1TN), and fibroblast cell lines (TD602 Rosie) were cultured in RPMI 1640 (Gibco^TM^, #11875093) supplemented with 10% heat-inactivated foetal bovine serum (CellSera, #AU-FBS/PG), 50 µM 2-mercaptoethanol (Sigma-Aldrich, #28713-500G-F), 10 mM HEPES (Gibco^TM^, #15630080), and 1% Antibiotic-Antimycotic (Gibco^TM^, #15240062) at 37 °C and 5% CO_2_ in a humidified incubator. Primary fibroblast cells were cultured with the same culture medium with added 10% AmnioMAX™-II Complete Medium (Gibco™, #11269016). Cells were harvested at 80% confluence for genomic DNA extraction using PureLink™ Genomic DNA Mini Kit (Invitrogen™, #K182001) according to the manufacturer’s protocol.

Solid biopsies and swabs were taken from deceased and live devils. For deceased devils, swab samples of the superficial layer of tissue sites suspected of DFT1 or DFT2 were taken. A 0.5 cm^3^ solid biopsy was taken from the same tissue site using a 4mm disposable biopsy punch (Kai Medical, #EUR0450/10). This was conducted as part of the routine necropsies performed at the Tasmanian Museum and Art Gallery (TMAG). Swab and biopsy samples were either flash frozen on dry ice for diagnosis in the laboratory or added into SHERLOCK parasite rapid extraction protocol (S-PREP buffer) for field diagnosis. S-PREP formulation was obtained from [28]. The buffer consists of nuclease free TE buffer (Invitrogen™, #AM9849) supplemented with 50 mM DTT and 20% (w/v) Chelex-100 (Bio-Rad, #1432832). Additionally, non-DFT solid biopsy samples from two devils were included. These were anal sac tumour and round cell tumour. For live devils with ulcerated lesions, swab samples were taken and/or a biopsy from the same spot using a 4 mm disposable biopsy punch (Kai Medical, #EUR0450/10). Samples were taken from the area with or without cleaning with saline and gauze. If the tissue sites suspected of DFT1 or DFT2 were not ulcerated, either a swab or fine needle aspirate (FNA) samples were taken. Swab, biopsy and FNA samples were added into S-PREP buffer for downstream SHERLOCK diagnosis. Additionally, ear biopsies (0.5 cm^3^) were collected from wild and quolls in a similar manner. **Supplementary Table 1** shows a list of live devils and quolls that the samples were collected from alongside their clinical presentation.

As working with endangered species can raise ethical concerns such as animal welfare and ecological impact, research must minimise harm through non-invasive methods whilst strictly adhering to ethical guidelines. This ensures that our work supports scientific advancement and species protection. Therefore, all samples were collected under the approval of the University of Tasmania Animal Ethics Committee (Permit Numbers: A0026159) and Department of Natural Resources and Environment (NRE) permit that protects threatened fauna used for scientific purposes (Permit Number: TFA22462).

### Overview of the SHERLOCK diagnostic assay for DFT1 and DFT2

Our SHERLOCK assay consists of 3 separate tubes. Tube 1 for crude DNA extraction, tube 2 for target DNA amplification, and tube 3 for the conversion of DNA to RNA and CRISPR-Cas13a nucleic acid detection (**Figure 1**). For tube 1, primary samples are incubated in S-PREP buffer at 95 °C for 10 minutes (**Figure 1A-1B**). The crude DNA sample from tube 1 is then added to tube 2 to amplify the target DNA with recombinase polymerase amplification (RPA) at 37 °C for 30 minutes (**Figure 1A**). RPA uses paired forward and reverse primers to amplify DFT1 or DFT2 specific markers, alongside primers specific to a housekeeping gene (RPL13a) which serves as a positive control (**Figure 1A**). The 5’ RPA primer also includes a T7 polymerase recognition site sequence, which is needed to convert DNA to RNA during the *in vitro* transcription (IVT) in tube 3 (**Figure 1A**). Tube 3 also contains the LwCas13a enzyme, the CRISPR-RNA (crRNA) specific to DFT1, DFT2, or RPL13, and quencher linked to fluorophore by a short RNA strand. Recognition of the target sequence by the crRNA activates the LwCas13a enzyme to cleave the RNA linking the quencher to the fluorophore. The conversion of DNA to RNA and detection of nucleic acid occurs simultaneously in tube 3 as it is incubated at 37 °C from 5 minutes to 3 hours, depending on the concentration of the target DNA (**Figure 1A**). Biopsies are taken from tumour site, or control site (**Figure 1B** shows an oral mucosal swab being performed). The test result is visualised using a handheld fluorescence viewer (minipcrbio^®^) equipped with blue LED lights (**Figure 1C**). All aforementioned steps can be performed in a field setting with limited instrumentation (**Figure 1D**).

**Figure 1.**
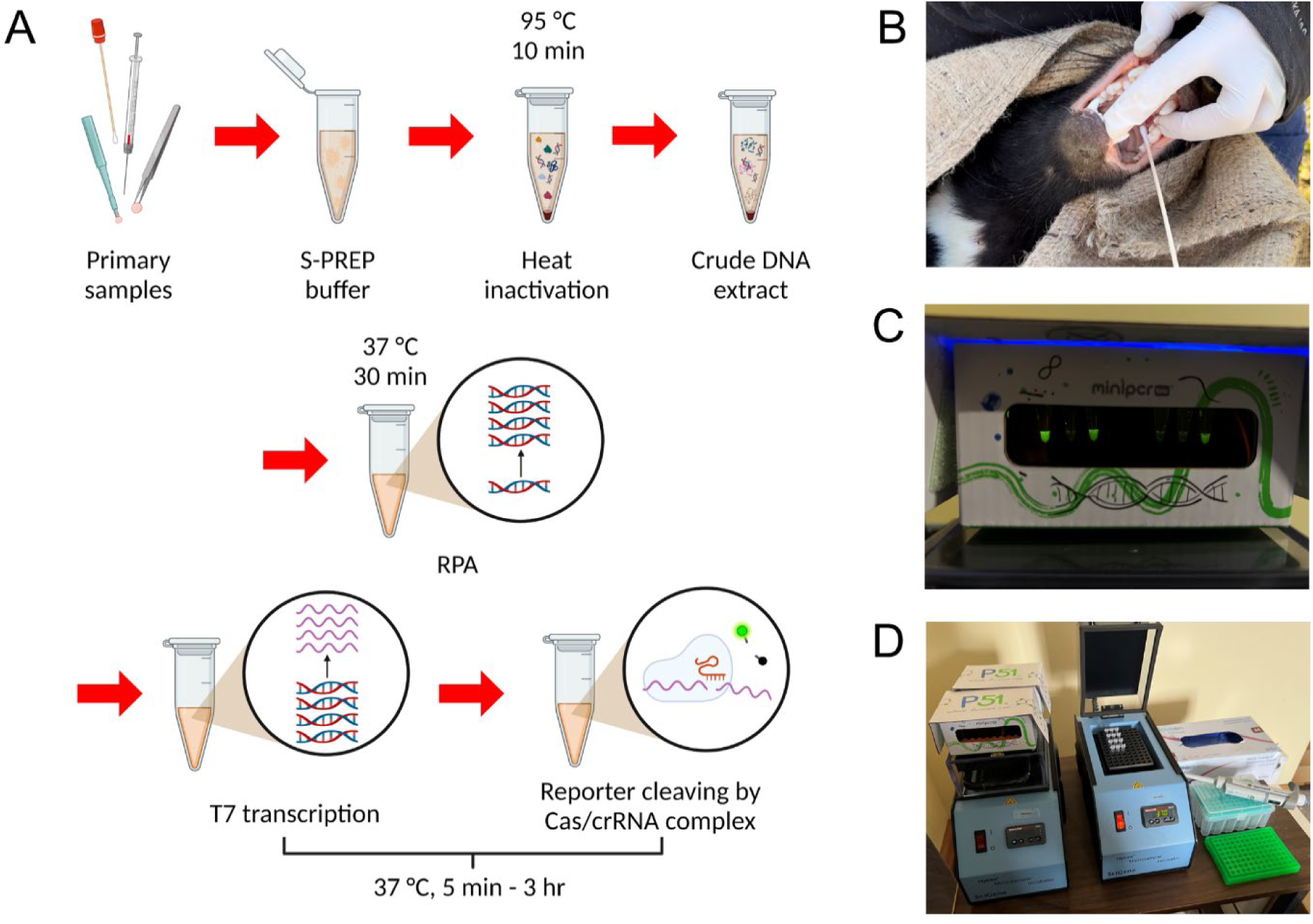
SHERLOCK field testing overview. A) Schematic of the SHERLOCK assay consisting of sample preparation involving the deactivation of primary samples in S-PREP buffer at 95 °C for 10 minutes, DNA amplification using RPA at 37 °C for 30 minutes, and T7 transcription and reporter cleaving at 37 °C for 5 minutes up to 3 hours. B) An oral mucosa swab sample was taken from a wild Tasmanian devil. C) SHERLOCK tests visualised using a handheld fluorescence viewer. D) Equipment needed to perform the field SHERLOCK diagnosis.

### Expression and purification of LwCas13a

Rosetta 2(DE3)pLysS Singles competent bacteria pre-transformed with pC013 -Twinstrep-SUMO-huLwCas13a (Addgene #90097) were used as an expression system for LwCas13a. A Luria-Bertani agar (LB) plate (Millipore, #28713-500G-F) containing 100 ug/mL of ampicillin was inoculated with bacteria and incubated overnight at 37 °C. Twenty-five mL of Terrific Broth (TB) medium (Invitrogen^TM^, #22711022) containing 100 µg/mL of ampicillin (Sigma-Aldrich, #A9518) was inoculated with a single bacteria colony and incubated overnight at 37 °C at 300 rpm. Four litres of TB medium containing 100 µg/mL ampicillin was inoculated with 20 mL of starter culture. Once the culture reached an optical density (OD) 0.4–0.6, the culture was allowed to cool to room temperature for 30 minutes. The expression of the gene encoding the Twinstrep-SUMO-huLwCas13a recombinant proteins was induced using 4 mL of 0.5 M Isopropyl ß-D-1-thiogalactopyranoside (IPTG, Sigma-Aldrich, #I6758-1G), and the culture was incubated overnight at 21 °C in a shaker at 200 rpm.

The bacteria were harvested by centrifugation at 5,200 × *g* for 15 minutes at 4 °C. Four mL of supplemented lysis buffer (20 mM Tris-HCl pH 8.0, 500 mM NaCl, 1 mM DTT, 1× cOmplete™ ULTRA protease inhibitor cocktail (Roche, #11873580001), 1 mg/mL lysozyme (Sigma-Aldrich, #L6876-1G), 1,500 units Benzonase (Millipore, #E1014-5KU)) per 1 g pelleted bacteria was used. The mixture was placed on the magnetic stirrer for 30 minutes at 4 °C before sonication on ice at the maximum amplitude with 1 s on and 2 s off for the total of 10 minutes of sonication time. The lysate was cleared by centrifugation at 10,000 rpm at 4 °C for 1 hour.

The clear fraction was decanted into a 250-ml tube and 5 ml Strep-Tactin Superflow plus (Qiagen, #30004) was added to batch bind the recombinant protein. The mixture was incubated for 2 hours at 4 °C with gentle shaking. A 100-mL glass chromatography column was prepared in the meantime by washing twice with cold lysis buffer (20 mM Tris-HCl pH 8.0, 500 mM NaCl, 1 mM DTT (Millipore, #3860-5GM)). Twenty mL of cold lysis buffer was then added to equilibrate the column and then drained immediately before addition of the resin solution. The column was then washed with 25 mL cold lysis buffer for 3 times. Fifteen mL of SUMO protease cleavage solution (lysis buffer, 250 units SUMO protease (Sigma-Aldrich, #SAE0067-2500UN), 22.5 μL TERGITOL™ NP-40 (Sigma-Aldrich, #NP40S-100ML) was added into the washed sample-resin. The SUMO protease cleavage was allowed to proceed overnight at 4 °C on the roller. The column was drained and washed 3 times with 5 mL cold lysis buffer to ensure the complete elution of untagged LwCas13a.

The collected sample was purified using ÄKTA Start protein purification system equipped with a 1 mL Sepharose High Performance (HiTrap SP HP) strong cation exchange column (Cytiva, #17115101) to remove SUMO protease, nucleases, and other unwanted contaminants. The column was prepared by washing with 5 column volumes of buffer B (20 mM Tris-HCl pH 7.5, 5% w/v glycerol (Sigma-Aldrich, #G5516-500ML), 1 mM DTT, 2 M NaCl), followed by 5 column volumes of buffer A (20 mM Tris-HCl pH 7.5, 5% w/v glycerol, 1 mM DTT) with a flow rate of 1 mL/min. The column was equilibrated in 12.5% buffer B (to achieve 250 mM NaCl) with a flow rate of 1 mL/min. The untagged LwCas13a was diluted by two-fold with buffer A to lower the NaCl concentration to 250 mM. The diluted sample was applied to the column, followed by column wash with 5 column volumes 12.5% buffer B, gradient elution with 10 column volumes from 12.5% buffer B to 100% buffer B.

Fractions containing LwCas13a were pooled and concentrated using 50 MWCO 15-mL spin filter (4,000 × *g* for 15 min). The buffer was exchanged by adding 15 mL protein storage buffer (50 mM Tris-HCl pH 7.5, 600 mM NaCl, 5% v/v glycerol, 2 mM DTT) to the same spin filter and centrifuged at 4,000 × *g* for 15 min at 4 °C. The purified LwCas13a was diluted to 63.3 µg/mL in protein storage buffer for use with CRISPR detection step. Protein fractions from each stage of purification were analysed using sodium dodecyl-sulfate polyacrylamide gel electrophoresis (SDS-PAGE) to ensure the successful expression and purification. Briefly, 4 μL 10X Bolt™ sample reducing agent (Thermo Fisher Scientific, #B0009), 10 µL 4X Bolt™ LDS Sample Buffer (Thermo Fisher Scientific, #B0007), and 16 µL UltraPure™ water (Invitrogen™, #10977015) were added to 10 µl of samples before heated at 95 °C for 5 minutes then incubated on ice for 2 minutes. Thirty µL of samples were loaded into Bolt™ 4-12% Bis-Tris Plus Mini Protein Gels (Thermo Fisher Scientific, #NW04120BOX) in 1x Bolt™ MES SDS running buffer (Thermo Fisher Scientific, #B0002). The gel was run at 200 V for approximately 20 minutes. The gel was stained for total protein using GelCode™ Blue Safe Protein Stain (Thermo Fisher Scientific, #24594) according to the manufacturer’s protocol before being visualised (**Supplementary Figure 1**).

### RPA primer design and screening

RPA primers were designed to amplify DFT1 and DFT2 markers, found on Chr2_GL841420:ChrX_GL867598 and Chr4_GL856969:Chr5_GL861630, respectively [10]. For DFT1, five forward primers and seven reverse primers were designed (**Supplementary Table 2**). For DFT2, five forward and five reverse primers were designed (**Supplementary Table 2**). Additionally, five forward and five reverse primers were designed for RPL13a to serve as an internal positive control (**Supplementary Table 2**). A region of the RPL13a sequence that was conserved between devil and brushtail possum (*Trichosurus vulpecula*) was selected, with the aim of developing RPA primers that could be used in other marsupial species.

All forward primers have a T7 RNA polymerase promoter sequence appended on the 5’ terminus (5’ – gaaattaatacgactcactataggg – 3’). Primers were designed as suggested by the manufacturer’s RPA guidelines (TwistAmp^®^ Basic Kit, TwistDx). RPA primers specific for ZIKV were also included as a positive control for RPA and downstream CRISPR detection reaction [30] (**Supplementary Table 2**). All primers were synthesised by Integrated DNA Technologies (IDT). For primer screening, products from all primer combinations were extracted using NucleoSpin Gel and PCR Clean-up kit (Macherey-Nagel, #740609.250), then run on a 3% agarose gel. Top candidates were selected for further screening by coupling with SHERLOCK and various crRNAs, as described below.

### RPA amplification

TwistAmp^®^ Basic Kit (TwistDx, # TABAS03KIT) was used for our primer screening. The reaction mix was prepared for each set of primers. 2.4 µL of each primer (10 µM starting concentration) was added into 29.5 µL of TwistAmp rehydration buffer. 8.65 µL of water was then added into the mixture before a quick vortex and spin down, which resulted in 42.95 µL total volume. 40 µL of the reaction mix was added directly into RPA pellet. 20 µL of the mixture was added into fresh PCR tubes (one reconstituted RPA pellet yields two RPA reactions). Unnecessary pipette tip changing was avoided as a significant amount of reaction volume could be lost due to the viscosity of the RPA reaction. 2 µL of the sample in S-PREP buffer or gDNA (10 ng/µL starting concentration for primer screening) were added into corresponding RPA reactions. 1.25 µL of magnesium acetate (MgOAc) provided with the TwistAmp^®^ Basic Kit (TwistDx) was then added to activate the reaction before the reaction was vortexed and centrifuged. The reaction was incubated at 37 °C for 30 minutes (this was done as soon as possible as the reaction starts the moment MgOAc is added). Post amplification product was directly used for CRISPR-LwCas13a detection step.

### S-PREP buffer preparation and crude DNA extraction

S-PREP formulation was obtained from [28]. The buffer consists of nuclease free TE buffer (Invitrogen™, #AM9849) supplemented with 50 mM DTT and 20% (w/v) Chelex-100 (Bio-Rad, #1432832). The strong chelating and reducing agents minimise the risk of enzymatic activity inhibition by inhibitors that are present in the crude lysate. Primary samples (from biopsy punches, fine needle aspirates, whole-blood samples, and swabs) were added to 300 µL S-PREP buffer in 1.5 ml microfuge tubes. Samples were then incubated at 95 °C for 10 minutes to inactivate enzymes and enzymatic inhibitors, and to release crude DNA from cells. The mixture was left to cool for two minutes on ice before being used in RPA amplification.

### CRISPR RNA (crRNA) design and screening

crRNAs were designed as suggested by Kellner, Koob [30]. One crRNA was designed for each DFT1 and DFT2 (**Supplementary Table 2**). ZIKV crRNA sequence was obtained from Kellner, Koob [30] (**Supplementary Table 2**). DFT1, DFT2, and ZIKV crRNAs were synthesised by IDT (Ultramer^TM^ RNA Oligo, 4 nmol, standard desalting). For RPL13a, three crRNA DNA templates were designed for *in vitro* transcription (IVT) (synthesised by IDT) (**Supplementary Table 2**). The annealing reaction for each crRNA was prepared by adding 1 μL crRNA DNA template (100 μM), 1 μL T7-3G oligonucleotide (100 μM), 1 μL Standard Taq buffer (10x) (NEB, #B9014S), and 7 μL UltraPure™ water into a PCR tube. The reaction was performed in a thermal cycler with a 5-minute denaturation at 95 °C and cooled to 4 °C using a 2.5% ramp rate. The annealed crRNA DNA templates were then added to the IVT reaction. For each crRNA, 10 μL annealed template was added into a PCR tube with 10 μL NTP buffer mix (HiScribe^®^ T7 Quick High Yield RNA Synthesis Kit, NEB, #E2040S), 2 μL T7 RNA Polymerase Mix (HiScribe^®^ T7 Quick High Yield RNA Synthesis Kit, NEB, #E2040S), and 17 μL UltraPure™ water. The reaction was incubated at 37 °C for 16 hours and then held at 4 °C prior to purification. crRNA was then purified using TRI Reagent^®^ (Sigma-Aldrich^®^, #T9424-25ML) according to the manufacturer’s protocol. The quality of crRNA was assessed using RNA ScreenTape for the 4200 TapeStation system (Agilent, #5067-5576). The best performing RPL13a crRNA was determined using SHERLOCK, which was then selected for high quality synthesis by IDT (Ultramer RNA Oligo™, 4 nmol, standard desalting).

For crRNA screening, DFT1, DFT2, RPL13a, and ZIKV templates were pre-amplified with RPA primers by PCR. DFT1 and DFT2 targets were amplified from DFT1 and DFT2 cell lines whilst RPL13a target was amplified from a devil primary fibroblast culture. ZIKV template was amplified from a short synthetic DNA fragment (IDT). The resulting target for IVT and CRISPR-Cas13a activation and cleaving (hereafter referred to as CRISPR reaction) contained a T7 sequence appended at the 5’ end. Ten ng of each template was used in the CRISPR reaction. In addition to the no template control (NTC), crRNA was mispaired to wrong targets to assess the specificity of the detection. The incubation and fluorescence reading were carried out at 37 °C for up to 3 hours using black 384-well flat, clear-bottom plates in a Spark^®^ 20M (Tecan) plate reader.

### CRISPR reaction mix composition and detection

The SHERLOCK ready mix was prepared in an RNase-free environment. For fluorescent plate viewing, the reaction was carried out in black 384-well flat, clear-bottom, black plate (Corning^®^). A Spark^®^ 20M (Tecan) plate reader was set to a kinetic fluorescence reading mode with 490 nm excitation (bandwidth of 10 nm) wavelength and 520 nm emission wavelength (bandwidth of 15 nm) with a manual gain setting of 100 (with gain regulation, 30 flashes, and 40 µs integration time). Z-position was set to automatically calculate from well A1. The stage temperature was set to 37 °C with one-minute measurement interval for one-hour or five-minute measurement interval over three hours. Each reaction (well) consists of 11.27 µL UltraPure™ water (Invitrogen^TM^), 0.4 µL HEPES buffer (1 M, pH adjusted to 6.8, Gibco), 0.18 µL MgCl_2_ (1M, Invitrogen^TM^), 0.8 µL ribonucleotide solution mix (NEB, #N0466S), 2 µL LwCas13a (63.3 µg/mL), 1 µL Murine RNase inhibitor (NEB, #M0314L), 0.5 µL T7 RNA polymerase (50,000 U/mL, NEB, #M0251S), 1 µL crRNA specific to the target of interest (10 ng/µL), 1.25 µL RNA reporter (2 µM starting concentration, 0.125 µM final concentration), and 1.6 µL post amplification RPA product. RNaseAlert™ Lab Test Kit v2 (Thermo Fisher Scientific) and RNaseAlert™ Substrate (IDT) were used as the RNA reporter. The reporter was reconstituted to 2 µM using 10× RNaseAlert buffer and nuclease-free water according to the manufacturer’s instructions prior to use.

For handheld fluorescence viewing amenable to field conditions, the reaction was carried out in 200-µL PCR tubes and incubated on a heat block or thermal cycler at 37 °C for up to three hours. Tubes were visualised using P51™ Molecular Fluorescence Viewer (minipcrbio^®^) equipped with blue LED strip and amber filter. Each CRISPR nucleic acid detection reaction consists of 9.52 µL UltraPure™ water, 0.4 µL 1M HEPES buffer, 0.18 µL 1M MgCl_2_ (1M, Invitrogen™, #AM9530G), 0.8 µL ribonucleotide solution mix, 2 µL LwCas13a (63.3 µg/mL), 1 µL Murine RNase inhibitor (NEB, #M0314L), 0.5 µL T7 RNA polymerase (50,000 U/mL, NEB, #M0251S), 1 µL crRNA specific to the target of interest (10 ng/µL), 3 µL RNA reporter (2 µM starting concentration, 0.3 µM final concentration), and 1.6 µL post amplification RPA product.

### Limit-of-detection

Limit-of-detection (LOD) was determined for SHERLOCK targeting DFT1, DFT2, and RPL13a. For each SHERLOCK assay, gDNA isolated from DFT1, DFT2, and fibroblast cells (for RPL13a) was serially diluted from 1 fM (10^-15^ M) down to 1 zM (10^-21^ M). Copy number was estimated using DNA Copy Number and Dilution Calculator (Thermo Fisher Scientific) based on the estimated Tasmanian devil’s genome size of 3 giga base pairs [32]. Two µL of the template from each concentration was added into the corresponding RPA reactions and incubated for 30 minutes. RPA products were then added into corresponding SHERLOCK ready mixes before being incubated in the Spark^®^ 20M (Tecan) plate reader with five-minute reading interval for three hours as previously described.

## RESULTS

### Screening of RPA primers, LwCas13a, and crRNA

RPA primers were screened to optimise primer pairs using RPA followed by gel electrophoresis. Six, three, and four RPA primer pairs for DFT1, DFT2, and RPL13a were selected based on the gel electrophoresis results. These primers were then tested with the sgRNA and LwCas13a in the SHERLOC reaction, which demonstrated that they efficiently amplified the target of interest leading to the CRISPR nucleic acid detection and reporter cleaving (**Supplementary Figures 2-4**).

The purified LwCas13a collateral cleaving ability was first tested by pairing crRNA to corresponding targets amplified by PCR, with a ZIKV DNA fragment as a positive control. The LwCas13a functioned as expected, with the fluorescence output rapidly increasing when pairing with ZIKV crRNA in the presence of ZIKV DNA fragment (**Figure 2A**). Additionally, the fluorescence kinetic showed that LwCas13a paired with our DFT1 and DFT2 crRNA led to a robust detection of the target of interest (**Figure 2A**). As the efficiency of DFT1 and DFT2 crRNA were comparable to the ZIKV crRNA previously developed and validated by Kellner, Koob [30], we did not perform any further crRNA screening. Mispairing DFT1 and DFT2 crRNAs and targets did not result in an activation of the CRISPR reaction as shown by the lack of fluorescence increase above background (**Figure 2A**). This suggested that each reaction is highly specific to the target of interest.

**Figure 2.**
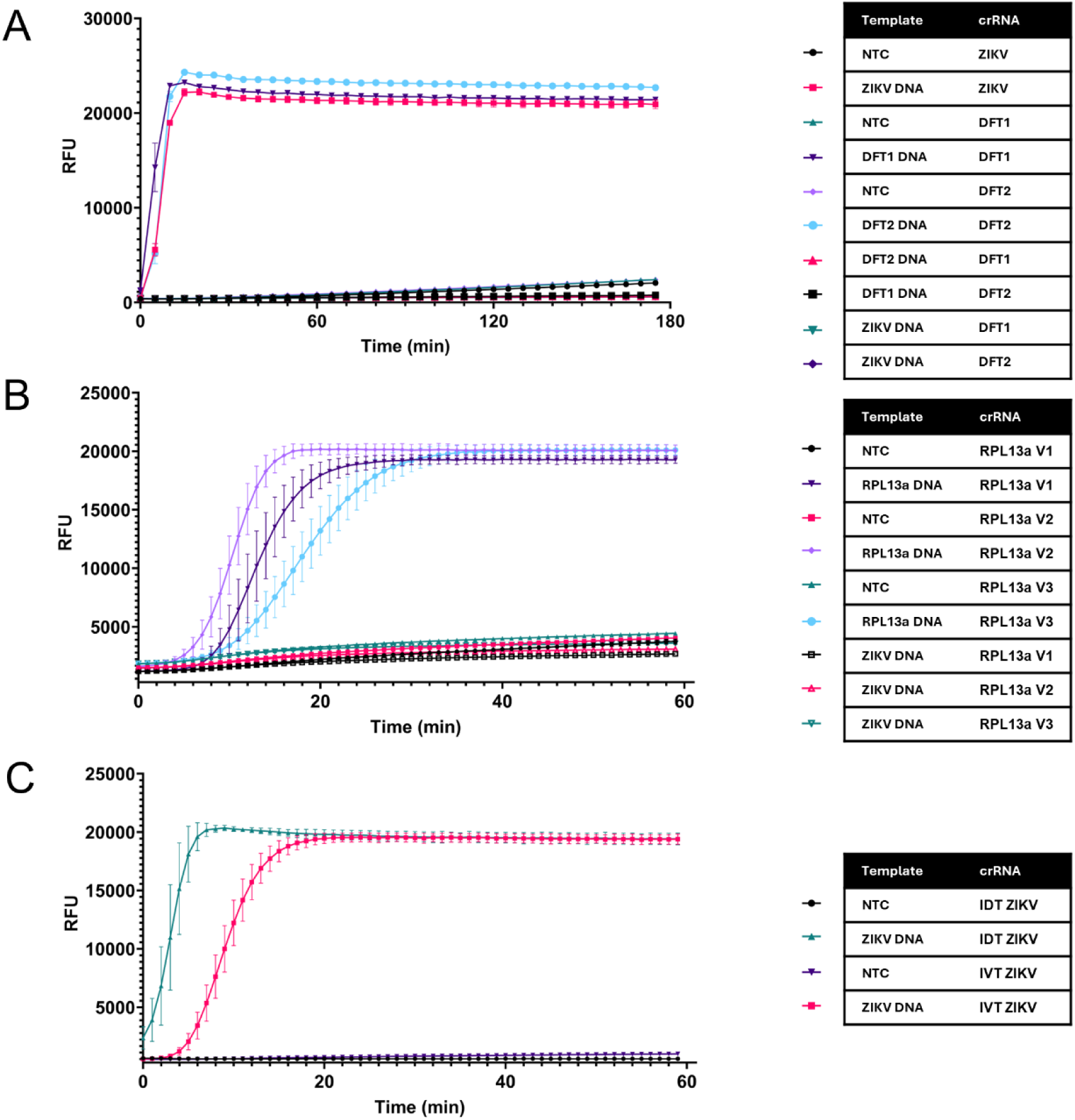
CrRNA specificity for the detection of DFT1, DFT2, and RPL13a. A) DFT1, DFT2, and ZIKV CRISPR reaction. crRNA was paired with the corresponding target, or water as a no template control (NTC). Additionally, crRNAs were paired with incorrect targets to assess the detection specificity. B) Screening of *in vitro* transcribed (IVT) RPL13a crRNAs. All IVT crRNAs were paired with their correct target, or water (NTC). The specificity of all IVT crRNAs were assessed by mispairing with ZIKV DNA. C) Assessing the efficiency between IVT crRNA and IDT synthesised crRNA. The crRNAs were either paired with their corresponding target or water (NTC). All reactions were assessed according to fluorescence kinetics of relative fluorescence unit (RFU) over time.

For RPL13a, we designed 3 crRNA DNA templates to be *in vitro* transcribed. The quality of all IVT crRNAs were assessed using RNA ScreenTape for the 4200 TapeStation system (Agilent). The result showed that all IVT reactions produced a sharp single RNA band with the correct size (**Supplementary Figure 5**). All three IVT RPL13a crRNAs produced detectable fluorescent signals when paired with their corresponding target, with the RPL13a V2 crRNA being the most efficient crRNA (**Figure 2B**). The specificity was again assessed by mispairing all crRNAs to the wrong target (ZIKV DNA in this case). The result suggested that all RPL13a crRNAs were highly specific for their target as indicated by the lack of fluorescence increase above background (**Figure 2B**). It was noted that the background fluorescence signal relative to the positive signal was higher for IVT crRNAs that commercially purchase crRNAs (**Figure 2B**). The efficiency between IVT crRNA and custom ordered crRNA (IDT) was additionally assessed. The commercially purchased crRNA had a higher detection efficiency than IVT crRNA despite the latter being of high quality and purity (**Figure 2C**). Therefore, we decided to custom order the RPL13a V2 crRNA to be able to achieve the highest sensitivity for our SHERLOCK tests.

### The effects of S-PREP buffer on downstream SHERLOCK detection

To assess the need for heating the samples in S-PREP buffer, the plasma samples from a healthy devil were spiked with DFT1 genomic DNA or DFT2 genomic DNA to simulate primary samples collected from DFT1- or DFT2-affected animals. The results revealed that SHERLOCK was able to detect its targets even without sample inactivation with S-PREP buffer at 95 °C (**Figure 3**). However, the rate of detection was markedly slower compared to S-PREP inactivated buffer (**Figure 3**). DFT1 and DFT2 SHERLOCK reached the maximum fluorescence signal after 180 minutes and 100 minutes of incubation, respectively, without S-PREP inactivation (**Figure 3A** and **3B**). The maximum fluorescence signal was achieved within 30 minutes of incubation with S-PREP inactivation (**Figure 3A** and **3B**).

**Figure 3.**
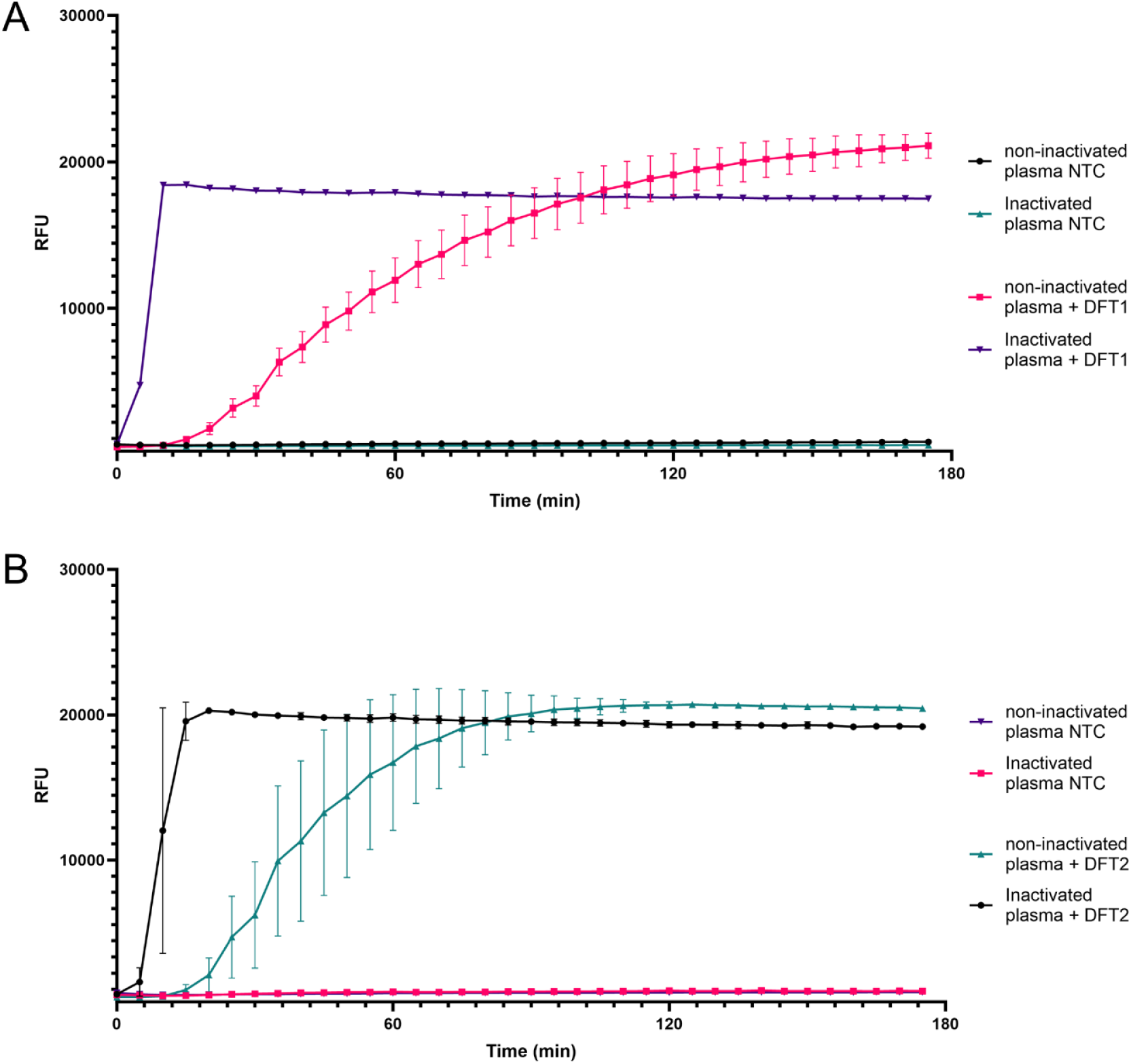
S-PREP vs no S-PREP for sample extraction/inactivation. Fluorescence kinetics (relative fluorescence unit, RFU) over time. A) Non-inactivated vs S-PREP inactivated plasma samples with or without spiked in DFT1 genomic DNA. B) Non-inactivated vs S-PREP inactivated plasma samples with or without spiked in DFT2 genomic DNA. NTC denotes no template control.

### SHERLOCK can distinguish DFT from non-DFT tumours

The ability to rule out non-DFT tumours is vital to monitor the spread of DFT1 and DFT2. We included two non-DFT tumour biopsies, an anal sac tumour and a round cell tumour, alongside ear biopsies from healthy devils, and performed a set of SHERLOCK tests following S-PREP inactivation and extraction. Results revealed that only when SHERLOCK was paired with the correct target, DFT1 or DFT2, it would lead to a positive detection indicated by the fluorescence increase (**Figure 4**). This further demonstrated the high level of specificity of our SHERLOCK assays when paired with primary sample biopsies (**Figure 4**).

**Figure 4.**
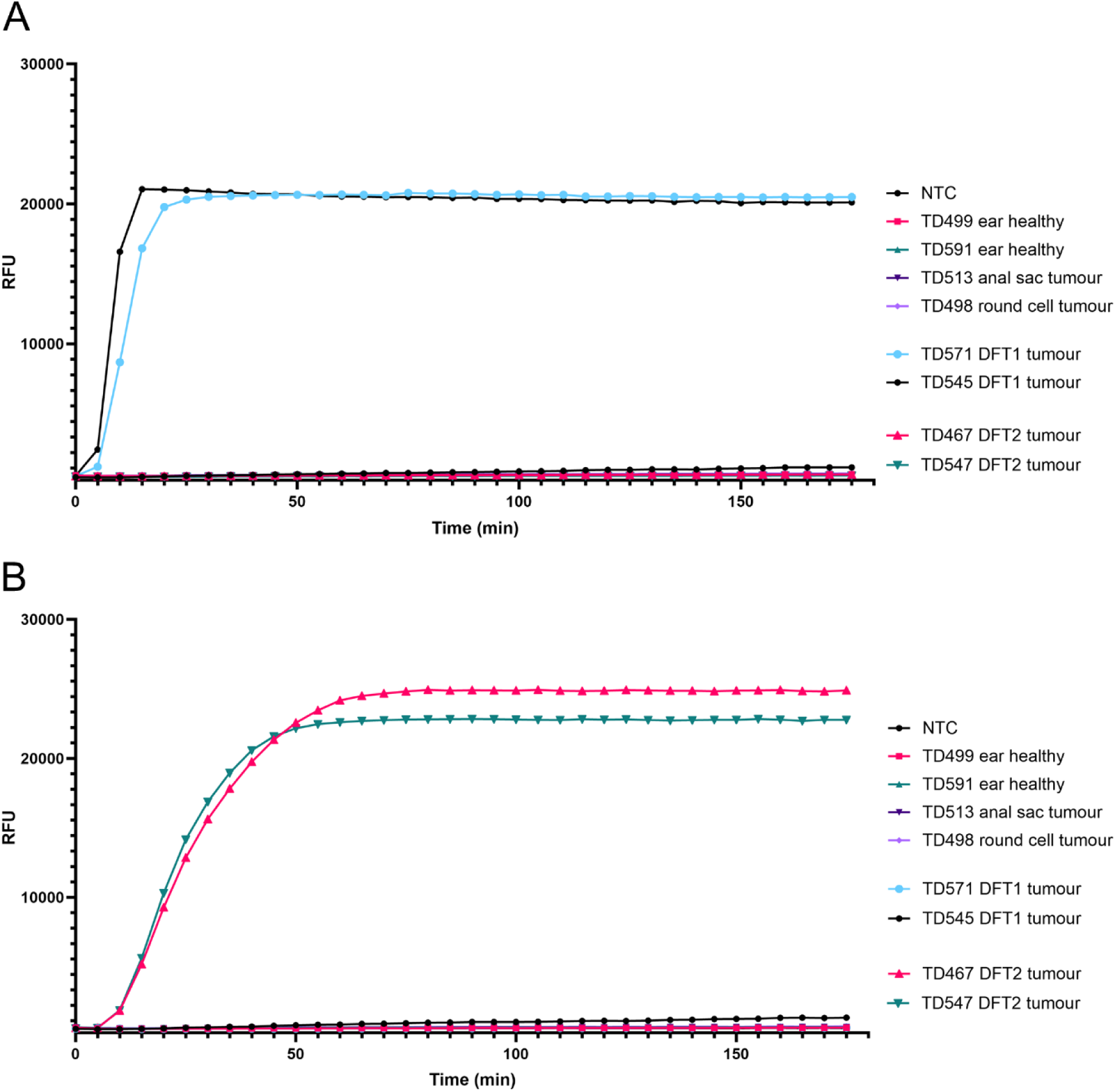
SHERLOCK on DFT1/2, non-DFT tumours, and healthy tissue biopsies. A) DFT1 SHERLOCK tested on DFT1 and DFT2, anal sac tumour, round cell tumour, and healthy ear biopsies. B) DFT2 SHERLOCK tested on DFT1 and DFT2, anal sac tumour, round cell tumour, and healthy ear biopsies. NTC and TD denote no template control and Tasmanian devil ID, respectively. Reactions were assessed as fluorescence kinetics of relative fluorescence unit, RFU) over time.

### SHERLOCK is highly sensitive

The sensitivity of SHERLOCK assays for DFT1, DFT2, and RPL13a was determined using serially diluted gDNA extracted from DFT1, DFT2, and fibroblast cells. We found that all three SHERLOCK assays could detect their respective target as low as 10 atto molar (aM) concentration, which is approximately the equivalent of 6 copies of DNA in 20 μL (**Figure 5**). No fluorescent signal higher than the background was detected in reactions below1 aM within 180 minutes of reaction time (**Figure 5**).

**Figure 5.**
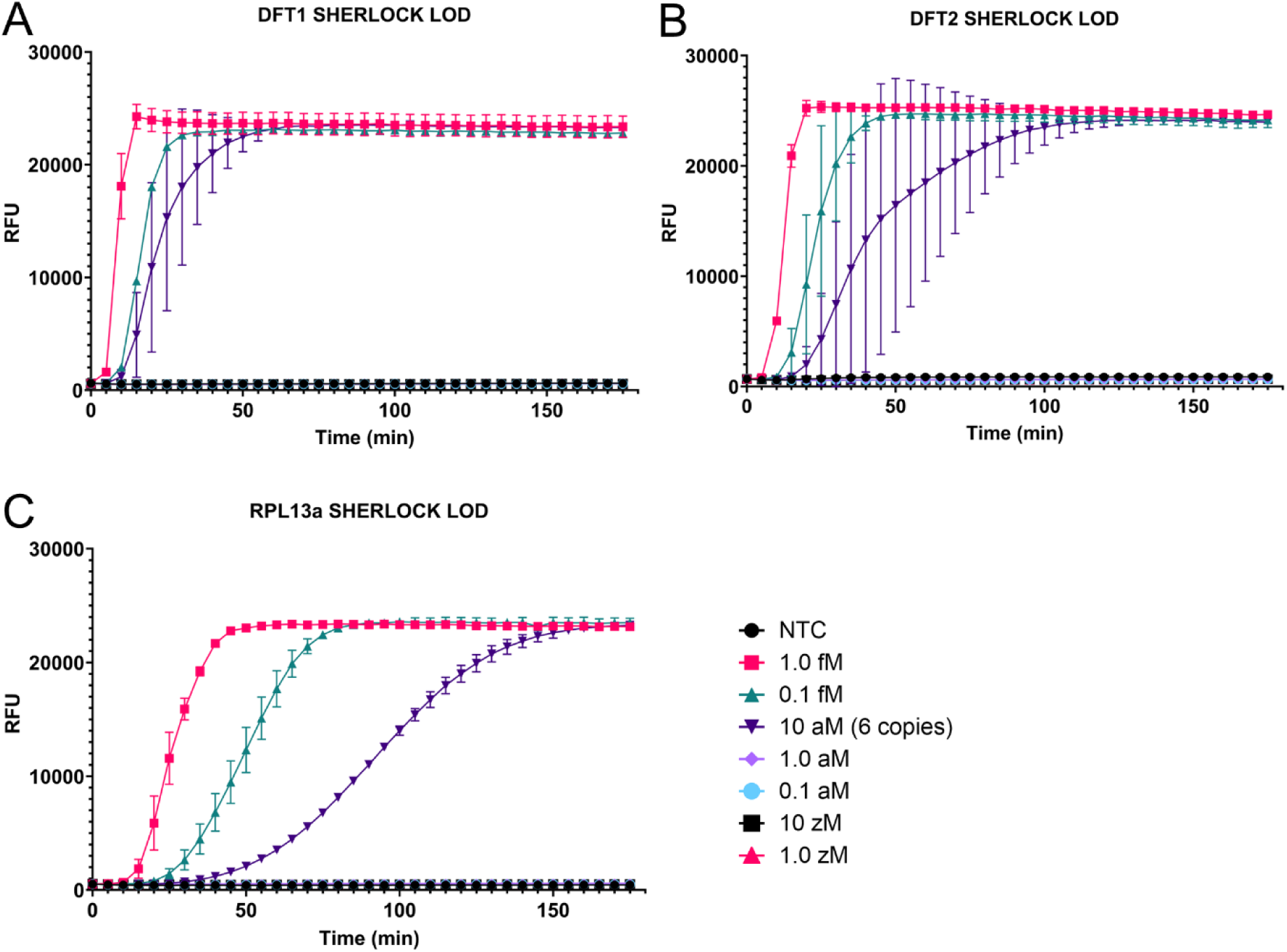
Limit-of-detection (LOD) of SHERLOCK for DFT1, DFT2, and RPL13a. Fluorescence kinetics (relative fluorescence unit, RFU) over time (minute, min) of three SHERLOCK assays. A serially diluted gDNA from DFT1, DFT2, and fibroblast cell lines from 1 femtomolar (fM) down to 1 zeptomolar (zM) was used to determine the LOD. A) SHERLOCK for DFT1, B) SHERLOCK for DFT2, and C) SHERLOCK for RPL13a. NTC denotes no template control.

### Optimisation and validation of field SHERLOCK testing on post-mortem samples

We validated our SHERLOCK DFT1 and DFT2 tests on post-mortem samples from deceased devils. Note that this was performed without the RPL13a control, which was not yet available at this time point. We performed SHERLOCK on biopsy and swab samples. Varying sizes of tumour biopsies were collected from six devils with suspected DFT1 and DFT2 masses before being added to microcentrifuge tubes with S-PREP buffer (**Figure 6A**). SHERLOCK tests with a biopsy of 5 mm^3^ or larger yielded a negative result whereas a biopsy smaller than 3 mm^3^ led to a positive detection of DFT1 and DFT2 in the first testing (**Figure 6A**). Subsequent testing was conducted on samples from devil TD618, TD619, and TD622 with 1:1 dilution in fresh S-PREP buffer. The false negative results were eliminated using the diluted crude lysate rather than bulky solid tissues **(Figure 6A**).

**Figure 6.**
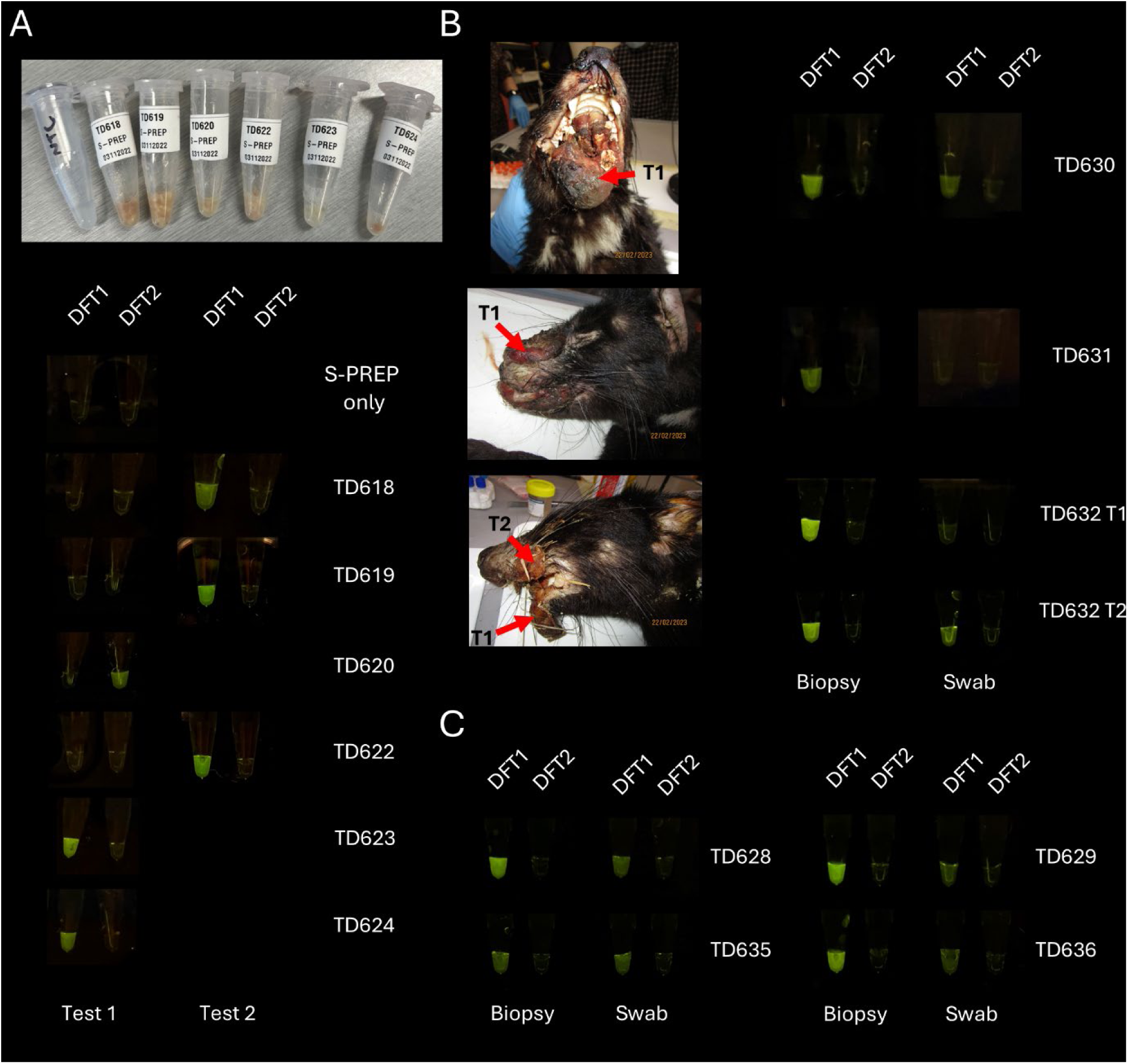
SHERLOCK testing on post-mortem samples. A positive detection for DFT1 or DFT2 is indicated by the visible green fluorescence. TD denotes Tamanian devil identification numbers. A) Tumour biopsy samples from six devils in S-PREP buffer and the corresponding SHERLOCK results. ‘S-PREP only’ indicates SHERLOCK on a no template control (NTC). Samples from TD618, TD619, and TD622 demonstrated a false-negative result that was eliminated when the sample was diluted 1:1 and re-tested. B) Pictures of three devils and corresponding SHERLOCK results from tumour biopsies and swab samples. Red arrows indicate where tumour samples were collected (T1 and T2 denotes tumour 1 and tumour 2, respectively). C) SHERLOCK on biopsy and swab samples from four devils. All biopsy and swab samples tested positive for DFT1.

Four swab and biopsy samples were taken from three deceased devils to assess the viability of SHERLOCK on swab samples (**Figure 6B**). Swab samples were taken from the area indicated on **Figure 6B** without cleaning the area prior to sample collection. Only two of four swab samples tested positive for DFT1 whilst all biopsy samples, the size of 1 mm^3^, produced a DFT1 positive detection (**Figure 6B**). Sample quality apart from the concentration of the DNA is an important consideration. We suspect that some samples, especially necrotic and/or with secondary microbial infection, can contain inhibitory compounds that can cause false negative results.

Lastly, we tested our SHERLOCK on four additional deceased animals to validate the optimisation conducted previously. Swab samples were taken from tissue regions that appeared to have less necrosis and microbial contamination. Biopsies of 1 mm^3^ were also collected to confirm the swab test results. **Figure 6C** shows that DFT1 was detected in all swab and biopsy samples for devils TD623, TD624, TD628, and TD635. The swabs tested positive but with lower fluorescence intensity than the biopsy samples at the time points used in this experiment (**Figure 6C**). **Table 1** summarises SHERLOCK test results from post-mortem samples.

**Table 1.** Result summary for SHERLOCK testing on post-mortem tumour samples.

|  | *Devil ID* | *Clinical presentation* | *PCR diagnosis* | *Sample type* | *DFT1 detection* | *DFT2 detection* |
| --- | --- | --- | --- | --- | --- | --- |
| *Figure 6A* | TD618 | DFTD | DFT1 | Biopsy | + | - |
|  | TD619 | DFTD | DFT1 | Biopsy | + | - |
|  | TD620 | DFTD | DFT2 | Biopsy | - | + |
|  | TD622 | DFTD | DFT1 | Biopsy | + | - |
|  | TD623 | DFTD | DFT1 | Biopsy | + | - |
|  | TD624 | DFTD | DFT1 | Biopsy | + | - |
| *Figure 6B* | TD630 | DFTD | DFT1 | Biopsy | + | - |
|  | TD630 | DFTD | DFT1 | Swab | + | - |
|  | TD631 | DFTD | DFT1 | Biopsy | + | - |
|  | TD631 | DFTD | DFT1 | Swab | - | - |
|  | TD632 T1 | DFTD | DFT1 | Biopsy | + | - |
|  | TD632 T1 | DFTD | DFT1 | Swab | - | - |
|  | TD632 T2 | DFTD | DFT1 | Biopsy | + | - |
|  | TD632 T2 | DFTD | DFT1 | Swab | + | - |
| *Figure 6C* | TD628 | DFTD | DFT1 | Biopsy | + | - |
|  | TD628 | DFTD | DFT1 | Swab | + | - |
|  | TD629 | DFTD | DFT1 | Biopsy | + | - |
|  | TD629 | DFTD | DFT1 | Swab | + | - |
|  | TD635 | DFTD | DFT1 | Biopsy | + | - |
|  | TD635 | DFTD | DFT1 | Swab | + | - |

\*, Positive detection achieved after the sample was diluted in fresh S-PREP buffer (1:1 dilution); 'T1' = tumour 1; 'T2' = tumour 2; '+' = positive result; '-' = negative result

### Point-of-care SHERLOCK test on wild animals in the field

We deployed the optimised DFT1, DFT2, and RPL13a SHERLOCK tests to assess performance in the field setting with minimally invasive swab samples and limited instrumentation. However, to demonstrate the versatility of the test, we also included biopsy and fine needle aspirate (FNA) samples. A total of 33 SHERLOCK tests were conducted on ten devils and three quolls. **Figure 7A** shows the complete SHERLOCK for DFT1, DFT2, and RPL13A positive control on wild devils with no clinical signs of DFTD (n = 3). The samples were positive for RPL13A, but no fluorescence signals were detected for DFT1 or DFT2 SHERLOCK, indicating that they were free of DFTD (**Figure 7A**). **Figure 7B** shows SHERLOCK with a positive detection of devils affected by DFTD (n = 5). The test was able to detect DFT1 infection in all devils with suspected DFT1 masses. However, two samples had to be re-tested due to a weak positive detection (Nacho) and a weak positive control detection in the FNA sample (Salsa) (**Figure 7B**). It was found that the weak fluorescence signal was likely due to the presence of inhibitors in the reaction. A swab sample from ‘Nacho’ was diluted in fresh S-PREP buffer (1:3 dilution) and retested, which revealed a stronger fluorescence intensity (**Figure 7B**). We added approximately 6 drops of FNA from Salsa into an S-PREP buffer which became apparent that the reaction was inhibited as indicated by a weak fluorescence signal. In a similar manner, an FNA sample from Salsa was diluted in fresh buffer (1:1 dilution), which also yielded a strong positive detection for DFT1 (**Figure 7B**).

**Figure 7.**
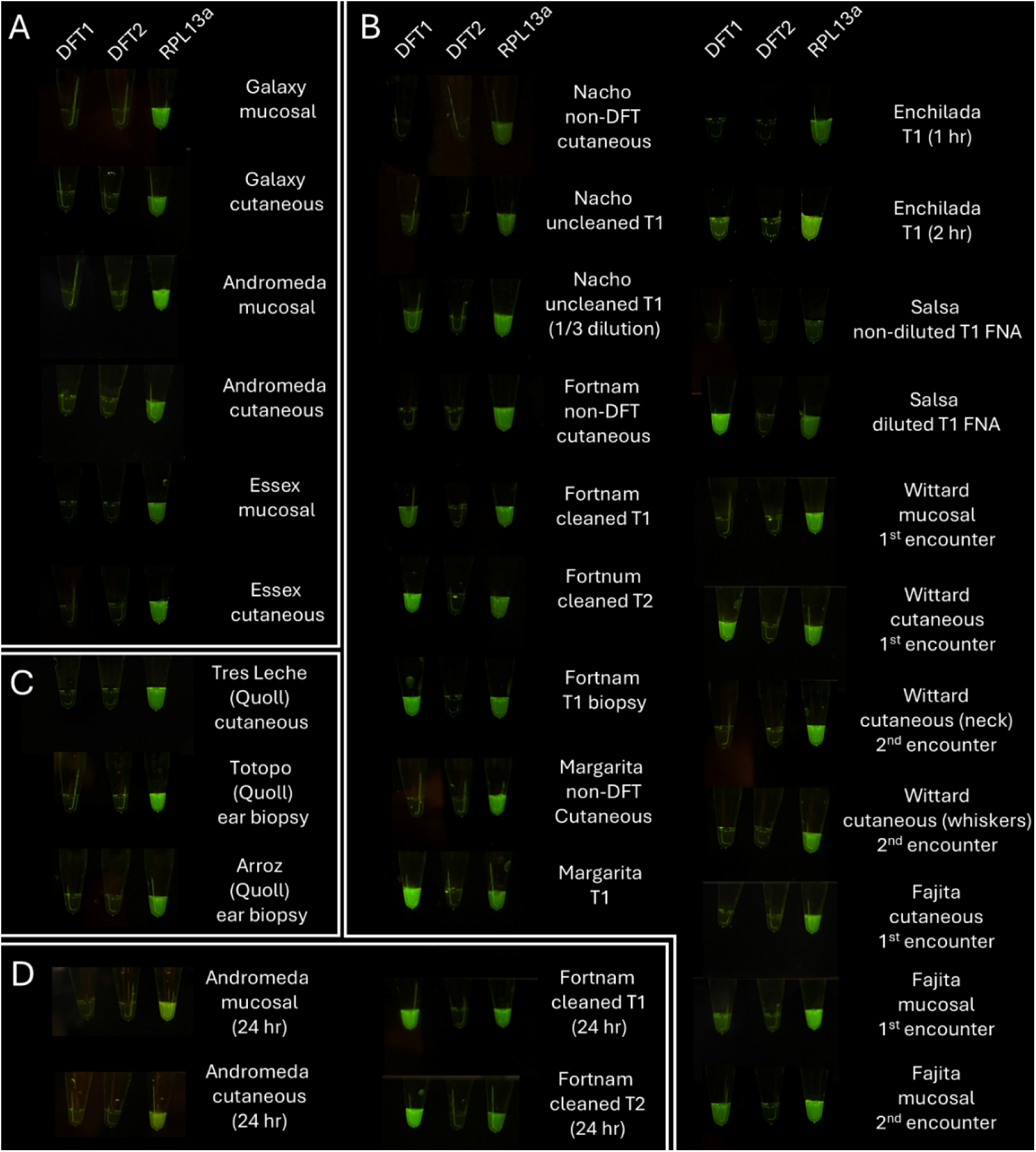
Field SHERLOCK testing on wild animals. Samples from cutaneous (cheek) and oral mucosal swabs were used unless stated otherwise. Positive detections were indicated by the visible green fluorescence. RPL13a served as a positive control. A) SHERLOCK results from DFT1 and DFT2 negative devils. B) SHERLOCK results from devils with DFT1. C) SHERLOCK results on spotted-tail quolls. All samples tested negative for DFT2 at this field site where DFT2 is not present. D) SHERLOCK results after 24 hours incubation. *T1,* tumour number 1*; T2,* tumour number 2*; uncleaned,* sample taken from an area without cleaning with saline and gauze; *cleaned,* sample taken from an area that has been cleaned with saline an gauze; *1^st^ encounter,* sample taken from an animal on the first encounter within the same trapping trip; *2^nd^ encounter,* sample taken from the same animal on a second encounter*; FNA,* fine needle aspirate.

To our surprise, two devils who appeared healthy and unaffected by DFTD, with no tumours or lesions, tested positive for DFT1. A cutaneous swab sample from Wittard, a four-year-old male devil, tested positive for DFT1 (**Figure 7B**). We repeated the test from samples taken on the same day to rule out cross contamination during sample preparation, which also resulted in DFT1 positive (data not shown). We collected more cutaneous samples from the same devil on the second trap encounter six days later, which revealed a negative result for DFT1 and DFT2 on both swab samples (**Figure 7B**). Another devil, Fajita, a one-year-old female devil with four pouch young that appeared healthy, tested positive for DFT1 from the mucosal swab (**Figure 7B**). Fajita was trapped again six days after the initial encounter and the mucosal swab sample again returned a DFT1 positive result (**Figure 7B**). Upon a closer inspection of the oral cavity, we identified a small non-ulcerated discoloured mark along the gum line (**Supplementary Figure 6**). However, this did not appear to have the clinical presentation resembling DFT1 or DFT2 lesion. All swabs taken from the lateral region away from any visible tumour (or area where DFT1 was detected in cases with no clinical sign of DFTD) returned negative results for DFT1 and DFT2 (n = 5). Additionally, the field was more than 200 km from regions known to have DFT2, and as expected all 33 samples tested negative for DFT2 at this site.

Lastly, we conducted SHERLOCK on swabs and biopsies taken from spotted-tail quolls (*Dasyurus maculatus*) to assess whether RPL13a can be detected and to help assess the false positive rate for the DFT1 and DFT2 tests. All swabs and biopsies tested positive for RPL13a including biopsies taken from spotted-tail quolls, suggesting that the RPL13a positive control test can work for multiple *Dasyuridae* species (**Figure 7C**). To test whether the fluorescence signal was observable after an extended period of incubation, DFT1 negative and positive samples from two devils were incubated for 24 hours. **Figure 7D** shows that the fluorescence signal remained visible after 24 hours and no positive detection was observed for reactions that were previously negative. This suggests that a longer incubation period is possible with no risk of false positive detection from the degradation of fluorescence reporter. **Table 2** summarises SHERLOCK POC tests results from live animals.

**Table 2.** Result summary for field SHERLOCK testing on wild animals.

|  | *Devil ID* | *Species* | *Clinical presentation* | *Sample type* | *Tissue type* | *Incubation time (hour)* | *DFT1 detection 1<sup>st</sup> encounter* | *DFT2 detection 1<sup>st</sup> encounter* | *RPL13a Detection 1<sup>st</sup> encounter* | *DFT1 detection 2<sup>nd</sup> encounter* | *DFT2 detection 2<sup>nd</sup> encounter* | *RPL13a Detection 2<sup>nd</sup> encounter* |
| --- | --- | --- | --- | --- | --- | --- | --- | --- | --- | --- | --- | --- |
| *Figure 7A* | Galaxy | *Sarcophilus harrisii* | Non-DFTD | Swab | Non-DFTD mucosal | 1 | - | - | + | NA | NA | NA |
|  | Galaxy | *Sarcophilus harrisii* | Non-DFTD | Swab | Non-DFTD cutaneous | 1 | - | - | + | NA | NA | NA |
|  | Andromeda | *Sarcophilus harrisii* | Non-DFTD | Swab | Non-DFTD mucosal | 1 | - | - | + | NA | NA | NA |
|  | Andromeda | *Sarcophilus harrisii* | Non-DFTD | Swab | Non-DFTD cutaneous | 1 | - | - | + | NA | NA | NA |
|  | Essex | *Sarcophilus harrisii* | Non-DFTD | Swab | Non-DFTD mucosal | 1 | - | - | + | NA | NA | NA |
|  | Essex | *Sarcophilus harrisii* | Non-DFTD | Swab | Non-DFTD cutaneous | 1 | - | - | + | NA | NA | NA |
| *Figure 7B* | Nacho | *Sarcophilus harrisii* | DFTD | Swab | Non-DFTD cutaneous | 1 | - | - | + | NA | NA | NA |
|  | Nacho | *Sarcophilus harrisii* | DFTD | Swab | Uncleaned T1 | 1 | + | - | + | NA | NA | NA |
|  | Nacho | *Sarcophilus harrisii* | DFTD | Swab | Uncleaned T1 (1/3 dilution) | 1 | + | - | + | NA | NA | NA |
|  | Fortnam | *Sarcophilus harrisii* | DFTD | Swab | Non-DFTD cutaneous | 1 | - | - | + | NA | NA | NA |
|  | Fortnam | *Sarcophilus harrisii* | DFTD | Swab | Cleaned T1 | 1 | + | - | + | NA | NA | NA |
|  | Fortnam | *Sarcophilus harrisii* | DFTD | Swab | Cleaned T2 | 1 | + | - | + | NA | NA | NA |
|  | Fortnam | *Sarcophilus harrisii* | DFTD | Biopsy | T1 | 1 | + | - | + | NA | NA | NA |
|  | Margarita | *Sarcophilus harrisii* | DFTD | Swab | Non-DFTD cutaneous | 1 | - | - | + | NA | NA | NA |
|  | Margarita | *Sarcophilus harrisii* | DFTD | Swab | T1 | 1 | + | - | + | NA | NA | NA |
|  | Enchilada | *Sarcophilus harrisii* | DFTD | Swab | T1 | 1 | - | - | + | NA | NA | NA |
|  | Enchilada | *Sarcophilus harrisii* | DFTD | Swab | T1 | 2 | + | - | + | NA | NA | NA |
|  | Salsa | *Sarcophilus harrisii* | DFTD | FNA | T1 | 1 | - | - | + | NA | NA | NA |
|  | Salsa | *Sarcophilus harrisii* | DFTD | FNA | T1 (1/1 dilution) | 1 | + | - | + | NA | NA | NA |
|  | Wittard | *Sarcophilus harrisii* | Non-DFTD | Swab | Non-DFTD mucosal | 1 | - | - | + | NA | NA | NA |
|  | Wittard | *Sarcophilus harrisii* | Non-DFTD | Swab | Non-DFTD cutaneous | 1 | + | - | + | NA | NA | NA |
|  | Wittard | *Sarcophilus harrisii* | Non-DFTD | Swab | Non-DFTD cutaneous (neck) | 1 | NA | NA | NA | - | - | + |
|  | Wittard | *Sarcophilus harrisii* | Non-DFTD | Swab | Non-DFTD cutaneous (whiskers) | 1 | NA | NA | NA | - | - | + |
|  | Fajita | *Sarcophilus harrisii* | Non-DFTD | Swab | Non-DFTD cutaneous | 1 | - | - | + | NA | NA | NA |
|  | Fajita | *Sarcophilus harrisii* | Non-DFTD | Swab | Non-DFTD mucosal | 1 | + | - | + | NA | NA | NA |
|  | Fajita | *Sarcophilus harrisii* | Non-DFTD | Swab | Non-DFTD mucosal | 1 | NA | NA | NA | + | - | + |
| *Figure 7C* | Tres Leche | *Dasyurus maculatus* | Non-DFTD | Swab | Non-DFTD cutaneous | 1 | + | - | + | NA | NA | NA |
|  | Totopo | *Dasyurus maculatus* | Non-DFTD | Biopsy | Non-DFTD ear | 1 | + | - | + | NA | NA | NA |
|  | Arroz | *Dasyurus maculatus* | Non-DFTD | Biopsy | Non-DFTD ear | 1 | + | - | + | NA | NA | NA |
| *Figure 7D* | Andromeda | *Sarcophilus harrisii* | Non-DFTD | Swab | Non-DFTD mucosal | 24 | - | - | + | NA | NA | NA |
|  | Andromeda | *Sarcophilus harrisii* | Non-DFTD | Swab | Non-DFTD cutaneous | 24 | - | - | + | NA | NA | NA |
|  | Fortnam | *Sarcophilus harrisii* | DFTD | Swab | Cleaned T1 | 24 | + | - | + | NA | NA | NA |
|  | Fortnam | *Sarcophilus harrisii* | DFTD | Swab | Cleaned T2 | 24 | + | - | + | NA | NA | NA |

‘\*’, low positive detection; ‘T1’, tumour 1; ‘T2’, tumour 2; ‘FNA’, fine needle aspirate; ‘+’, positive result; ‘-’, negative result; ‘NA’, not applicable

## DISCUSSION

SHERLOCK results could generally be obtained in less than an hour post-sample processing, providing a more rapid less invasive diagnostic method compared to other techniques such as PCR, immunohistochemistry, and mass spectrometry. PCR can be used with swabs or biopsies, but generally requires DNA extraction prior to amplification. This results in a longer time to result and increases the chance of sample contamination, particularly in the field. Immunohistochemistry requires the collection of solid biopsies, which is not always feasible if the tumour size is below a certain size. Additionally, solid biopsy collection via punch biopsy carries the risk of tumour spread to surrounding tissues and pushing microbes into subcutaneous regions. Mass spectrometry, used to detect proteins upregulated in DFT1 and DFT2, requires serum samples, which must be obtained under anaesthesia -introducing potential adverse effects on the animal and broader ecological consequences. As SHERLOCK is less invasive to other methods, this will allow us to minimise harm whilst strictly adhering to ethical guidelines. This ensures that our work minimises stress to a wild endangered species.

Our study demonstrated that the inactivation of samples in S-PREP buffer followed by SHERLOCK can reliably detect DFT1 and DFT2 with a high level of sensitivity and a low false positive rate in the laboratory and the field. Laboratory tests showed that as few as six copies of DNA per 20 μL reaction could be detected. The sensitivity of our DFT1, DFT2, and RPL13a SHERLOCK assays is on par with the SHERLOCK test for SARS-CoV-2 kit that gained the emergency use authorisation from the FDA (LOD of 6.75 copies of genomic viral RNA) [33]. The field results supported the sensitivity of the tests with all devils with suspected DFT1 tested positive for DFT1. Equally important was the low false positive rate, demonstrated by DFT2 testing negative in all 33 separate SHERLOCK tests across in regions where DFT2 is not present. Additionally, only devils are known to be infected with DFT1 and DFT2 cells, and all 3 quolls were negative for DFT1 and DFT2 but positive for RPL13a. This suggests that positive results are likely true positives if human error can be ruled out.

Two wild devils, Wittard and Fajita, that appeared to be healthy tested positive for DFT1. Both Wittard and Fajita were re-trapped six days after the initial positive DFT1 test. Fajita again tested positive for DFT1, and upon further inspection, a small non-ulcerated discolouration was detected in the oral cavity. The appearance of the discolouration does not align with what would typically be seen in small DFT1 masses. There is limited data on the early visual indicators of DFT1 due to the lack of rapid point-of-care tests. Whilst contamination of the trap or equipment is possible, it is unlikely, as none of DFT2 samples were positive and DFT1 had no false positives in samples known to be negative for DFT1. If contamination were the case, it would raise the question of why only this devil tested positive twice, whilst other devils that were re-trapped tested negative on multiple occasions. Additionally, SHERLOCK tests performed from the dorsal ventral region of the neck away from visible tumour in affected devils did not test positive despite the close proximity of the tumour, which makes it a less likely explanation. If future positive samples from animals with no visible tumours proves reliable and reanalysis of prior samples via Tasman PCR or other diagnostic methods show that the results were true positives, then it suggests that SHERLOCK could provide the crucial pre-clinical test that is needed to improve devil conservation management.

The strong positive sample from Wittard that was followed by a clear negative cannot be explained with the data available at this time. A devil with severe DFT1, Nacho, was trapped within 100 metres from Wittard. It is conceivable that Wittard interacted with Nacho before entering the trap, which could have resulted in residual DFT1 DNA on Wittard that led to the positive DFT1 test. However, contamination or false positive results remain possible explanations.

The minimal instrumentation required for SHERLOCK makes this the most feasible field diagnostic test to date. Additionally, the protocol is relatively simple compared to other techniques such as PCR. Thus, SHERLOCK tests for other pathogens could be used for other wildlife studies and in veterinary facilities that lack molecular biology equipment. Additional studies are needed to compare the sensitivity and specificity of alternative diagnostic methods. We found that sample extraction and inactivation with S-PREP is needed to extract crude gDNA and to prevent the inhibition of RPA and downstream CRISPR detection. This was observed when comparing S-PREP inactivated and non-inactivated plasma sample with spiked in DFT1 or DFT2 gDNA. Additionally, a study led by Lee, Puig [28] reported that non-inactivated samples could also result in a false positive detection. This was the case with samples with high level of nucleases, causing unspecific cleavage of the RNA reporter. As a rapid test is highly desirable, future studies can examine the minimum time needed for incubating and heating in SPREP buffer.

Our field testing demonstrated the versatility of SHERLOCK, which is compatible with various types of samples such as solid/liquid biopsy and swab sample. For primary biopsies, it was evident that too much starting sample could lead to an inhibition of SHERLOCK, likely due to the excess inhibitory molecules. This was eliminated when less sample was used or when the inactivated sample was diluted with fresh S-PREP buffer. We discovered that swab and FNA samples with a secondary infection are prone to giving a false negative result. Therefore, samples should be taken from area with less necrosis whenever possible.

The site of field testing does not have any DFT2 cases. Therefore, we could not compare the efficiency of DFT2 SHERLOCK test. However, samples from deceased animals suggested that the test can detect DFT2 in the simulated field setting with limited instrumentation. Together with current ongoing management efforts, our test will pave the way towards active management approaches that are not currently possible. For example, several therapeutic and vaccine approaches have been tested or are in development, and a devil that tested positive in the field for early-stage tumours could be administered a vaccine or therapy in the field [34–37].

Future studies should directly compare the specificity and sensitivity of the Tasman PCR test and SHERLOCK test for each sample collected. In the first case, the two tests could be run using aliquots from the same swab sample. Other diagnostic methods such as Oxford Nanopore sequencing may be used as an alternative to SHERLOCK or to validate SHERLOCK finding. However, the technique requires multistep preparation compared to SHERLOCK. Previous studies have showed that mass spectrometry could be used to identify DFT1- and DFT2-associated proteins in serum samples, and has the potential to detect pre-clinical DFTD [38, 39].

SHERLOCK testing on animals without clinical signs of DFTD, followed by subsequent trapping trips, will be a key validation step for the crucial pre-clinical test. If a devil without visible DFTD symptoms tests positive on SHERLOCK and later develops a facial tumour, this could indicate successful pre-clinical detection. Future studies should aim to establish the latent period of DFTD, provided positive results from asymptomatic animals prove reliable. Disease monitoring is essential for future vaccination trials. A subset of animals will be challenged with live tumour cells after vaccination. SHERLOCK could be employed to more precisely identify when tumour DNA can be detected and when clinical signs are observed. Sampling from animals regardless of their clinical status can help reduce potential selection bias.

Although the goal of this study was to assess the performance of SHERLOCK diagnostics, future work needs to explore the long-term scalability and real-world implementation. “One-pot” SHERLOCK assays could be developed to simplify the testing procedure and to reduce the risk of cross-contamination associated with opening and closing multiple tubes. This would also make the test more amenable to non-laboratory scientists. Furthermore, a smartphone application can be developed to be able to quantify the fluorescence signal from a handheld fluorescence viewer. To extend shelf life, they can potentially be lyophilised which would allow the test to be stored at room temperature. Lyophilisation of the reagents can also potentially simplify the test protocol and reduce the time required to prepare reagents for field testing. Various studies showed that lyophilisation is possible, and the same level of sensitivity is retained [28, 40, 41].

## Supporting information

Supplementary Table 1 - list of samples

Supplementary Table 2 - list of RPA primers and crRNAs

## ACKNOWLEDGEMENTS

Thank you to the field teams of the Save the Tasmanian Devil Program and the Department of Natural Resources and Environment Tasmania (NRE) Jodie Elmer, Sarah Peck, Judy Clarke, and Bill Brown who assisted with field sample collection and necropsy. We thank Jocelyn Darby for her ongoing contributions to the team. Thank you to Liz Murchison and her team at the Transmissible Cancer Group (Cambridge, UK), their genomic analysis provided the foundation for this study.

## AUTHOR CONTRIBUTIONS

Field sample collection: J.M.D.

Laboratory assay and analysis: A.K., R.K.C., D.A.G., J.M.D., W.Z.

Data analysis: A.K., R.K.C., D.A.G., J.M.D., W.Z.

Funding acquisition; A.S.F., A.K.

Writing – original draft: A.K.

Writing – review and editing: A.K., R.K.C., D.A.G., J.M.D., W.Z., C.E.B., K.A.F., A.S.F.

## REFERENCES

1. Hawkins, C.E., et al., Emerging disease and population decline of an island endemic, the Tasmanian devil Sarcophilus harrisii. Biological Conservation, 2006. 131(2): p. 307–324.

2. Brüniche–Olsen, A., et al., Ancient DNA tracks the mainland extinction and island survival of the Tasmanian devil. Journal of Biogeography, 2018. 45(5): p. 963–976.

3. White, L.C., et al., High-quality fossil dates support a synchronous, Late Holocene extinction of devils and thylacines in mainland Australia. Biology Letters, 2018. 14(1): p. 20170642.

4. Jones, M.E. and L.A. Barmuta, Niche Differentiation Among Sympatric Australian Dasyurid Carnivores. Journal of Mammalogy, 2000. 81(2): p. 434–447.

5. Guiler, E., Observations on the Tasmanian Devil, Sarcophilus harrisi (Dasyuridae: fvlarsupiala) at Granville Harbour, 1966-75. 1978.

6. Comte, S., et al., Changes in spatial organization following an acute epizootic: Tasmanian devils and their transmissible cancer. Global Ecology and Conservation, 2020. 22: p. e00993.

7. Lazenby, B.T., et al., Density trends and demographic signals uncover the long-term impact of transmissible cancer in Tasmanian devils. J Appl Ecol, 2018. 55(3): p. 1368–1379.

8. Pye, R.J., et al., A second transmissible cancer in Tasmanian devils. Proceedings of the National Academy of Sciences, 2016. 113(2): p. 374.

9. Stammnitz, M.R., et al., The Origins and Vulnerabilities of Two Transmissible Cancers in Tasmanian Devils. Cancer Cell, 2018. 33(4): p. 607–619.e15.

10. Kwon, Y.M., et al., Tasman-PCR: a genetic diagnostic assay for Tasmanian devil facial tumour diseases. Royal Society Open Science, 2018. 5(10): p. 180870.

11. Margres, M.J., et al., Spontaneous Tumor Regression in Tasmanian Devils Associated with RASL11A Activation. Genetics, 2020. 215(4): p. 1143–1152.

12. Pye, R., et al., Demonstration of immune responses against devil facial tumour disease in wild Tasmanian devils. Biology Letters, 2016. 12(10).

13. Wright, B., et al., Variants in the host genome may inhibit tumour growth in devil facial tumours: evidence from genome-wide association. Scientific Reports, 2017. 7(1): p. 423.

14. Kaminski, M.M., et al., CRISPR-based diagnostics. Nature Biomedical Engineering, 2021. 5(7): p. 643–656.

15. Notomi, T., et al., Loop-mediated isothermal amplification of DNA. Nucleic Acids Res, 2000. 28(12): p. E63.

16. Piepenburg, O., et al., DNA Detection Using Recombination Proteins. PLoS Biology, 2006. 4(7): p. e204.

17. Yan, S., et al., Comparison of four isothermal amplification techniques: LAMP, SEA, CPA, and RPA for the identification of chicken adulteration. Food Control, 2024. 159: p. 110302.

18. Pang, J., et al., A real-time recombinase polymerase amplification assay for the rapid detection of Vibrio harveyi. Molecular and Cellular Probes, 2019. 44: p. 8–13.

19. Cabada, M.M., et al., Recombinase Polymerase Amplification Compared to Real-Time Polymerase Chain Reaction Test for the Detection of Fasciola hepatica in Human Stool. Am J Trop Med Hyg, 2017. 96(2): p. 341–346.

20. Zou, Y., M.G. Mason, and J.R. Botella, Evaluation and improvement of isothermal amplification methods for point-of-need plant disease diagnostics. PLOS ONE, 2020. 15(6): p. e0235216.

21. Mojica, F.J.M., et al., Intervening Sequences of Regularly Spaced Prokaryotic Repeats Derive from Foreign Genetic Elements. Journal of Molecular Evolution, 2005. 60(2): p. 174–182.

22. Pourcel, C., G. Salvignol, and G. Vergnaud, CRISPR elements in Yersinia pestis acquire new repeats by preferential uptake of bacteriophage DNA, and provide additional tools for evolutionary studies. Microbiology, 2005. 151(3): p. 653–663.

23. Mali, P., et al., RNA-Guided Human Genome Engineering via Cas9. Science, 2013. 339(6121): p. 823–826.

24. Cong, L., et al., Multiplex Genome Engineering Using CRISPR/Cas Systems. Science, 2013. 339(6121): p. 819–823.

25. Chen, J.S., et al., CRISPR-Cas12a target binding unleashes indiscriminate single-stranded DNase activity. Science, 2018. 360(6387): p. 436–439.

26. Gootenberg, J.S., et al., Nucleic acid detection with CRISPR-Cas13a/C2c2. Science, 2017. 356(6336): p. 438–442.

27. Patchsung, M., et al., Clinical validation of a Cas13-based assay for the detection of SARS-CoV-2 RNA. Nature Biomedical Engineering, 2020. 4(12): p. 1140–1149.

28. Lee, R.A., et al., Ultrasensitive CRISPR-based diagnostic for field-applicable detection ofPlasmodiumspecies in symptomatic and asymptomatic malaria. Proceedings of the National Academy of Sciences, 2020. 117(41): p. 25722–25731.

29. Zahra, A., et al., The SHERLOCK Platform: An Insight into Advances in Viral Disease Diagnosis. Molecular Biotechnology, 2023. 65(5): p. 699–714.

30. Kellner, M.J., et al., SHERLOCK: nucleic acid detection with CRISPR nucleases. Nature Protocols, 2019. 14(10): p. 2986–3012.

31. Siddle, H.V., et al., Reversible epigenetic down-regulation of MHC molecules by devil facial tumour disease illustrates immune escape by a contagious cancer. Proceedings of the National Academy of Sciences, 2013. 110(13): p. 5103–5108.

32. Murchison, E.P., et al., Genome sequencing and analysis of the Tasmanian devil and its transmissible cancer. Cell, 2012. 148(4): p. 780–91.

33. BIOSCIENCES, S., Sherlock CRISPR SARS-CoV-2 kit. 2022, FDA.

34. Schönbichler, A., et al., Tyrosine kinase targeting uncovers oncogenic pathway plasticity in Tasmanian devil transmissible cancers. The EMBO Journal, 2025.

35. Maskell, K.G., et al., Differentially expressed growth factors and cytokines drive phenotypic changes in transmissible cancers. Discovery Immunology, 2025. 4(1).

36. Kayigwe, A.N., et al., A human adenovirus encoding IFN-γ can transduce Tasmanian devil facial tumour cells and upregulate MHC-I. Journal of General Virology, 2022. 103(11).

37. Flies, A.S., et al., An oral bait vaccination approach for the Tasmanian devil facial tumor diseases. Expert Review of Vaccines, 2020. 19(1): p. 1–10.

38. Espejo, C., et al., Extracellular vesicle proteomes of two transmissible cancers of Tasmanian devils reveal tenascin-C as a serum-based differential diagnostic biomarker. Cellular and Molecular Life Sciences, 2021. 78(23): p. 7537–7555.

39. Espejo, C., et al., Cathelicidin-3 Associated With Serum Extracellular Vesicles Enables Early Diagnosis of a Transmissible Cancer. Frontiers in Immunology, 2022. 13.

40. Li, J., et al., Optimizing two-step and one-step SHERLOCK nucleic acid detection assays and their applications in detecting influenza A and B viruses. Microchemical Journal, 2025. 211: p. 113064.

41. Puig, H.d., et al., Minimally instrumented SHERLOCK (miSHERLOCK) for CRISPR-based point-of-care diagnosis of SARS-CoV-2 and emerging variants. Science Advances, 2021. 7(32): p. eabh2944.

