## Supplementary Figures 1-6 for "Development of a point-of-care field diagnostic test for DFT1 and DFT2"

Development of a point-of-care field diagnostic test for DFT1 and DFT2

**AUTHORS**

Anuk Kruawan[1], Rajendra K.C.[2], David A Gell[1,2], Jocelyn M. Darby[1], Weizhen Zhu[1], Chrissie E. B. Ong[1], Kirsten A. Fairfax[2], Andrew S. Flies[1]*

**AFFILIATIONS**

[1]Menzies Institute for Medical Research, College of Health and Medicine, University of Tasmania, Hobart, TAS 7000, Australia

[2]School of Medicine, College of Health and Medicine, University of Tasmania, Hobart, TAS 7000, Australia

**CORRESPONDING AUTHOR CONTACT INFORMATION**

Menzies Institute for Medical Research, University of Tasmania

GPO Box 341

Hobart, Tasmania, 7001, Australiat: +61 3 6226 4614

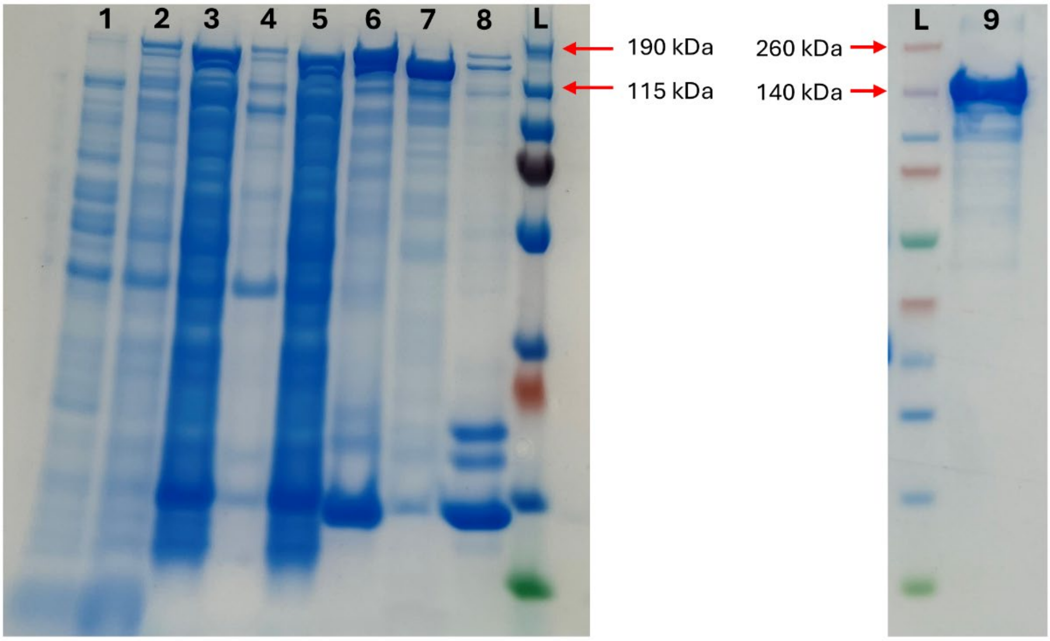


**Supplementary Figure 1. SDS-PAGE analysis for LwCas13a expression and purification.**

SDS-PAGE analysis of the progress of LwCas13a protein purification. The fractions are L, ladder; 1, uninduced bacterial culture; 2, IPTG induced bacterial culture; 3, cleared lysate; 4, pelleted lysate; 5, Strep-Tactin resin flowthrough non-binder; 6, Strep-Tactin resin-bound protein; 7, SUMO-cleaved protein; 8, Strep-Tactin resin after SUMO cleaving; 9, concentrated protein after cation exchange chromatography. LwCas13a protein is 138.51 kDa in size.

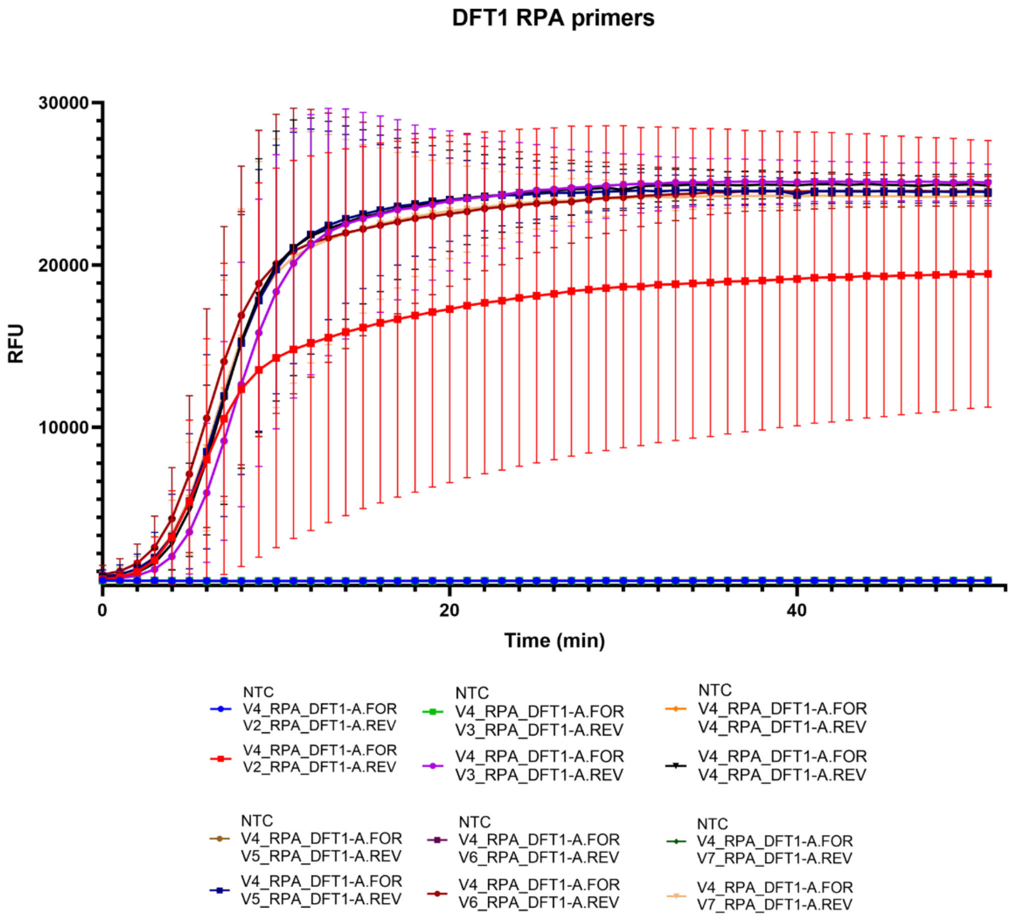


**Supplementary Figure 2. Fluorescence kinetics of top performing DFT1 RPA primers.**

Fluorescence kinetics (relative fluorescence unit, RFU) over time (minute, min). Six pairs of RPA primers for DFT1 were assessed to identify the best performing primer combination. These primers were paired with DFT1 gDNA or water as a no template control (NTC).

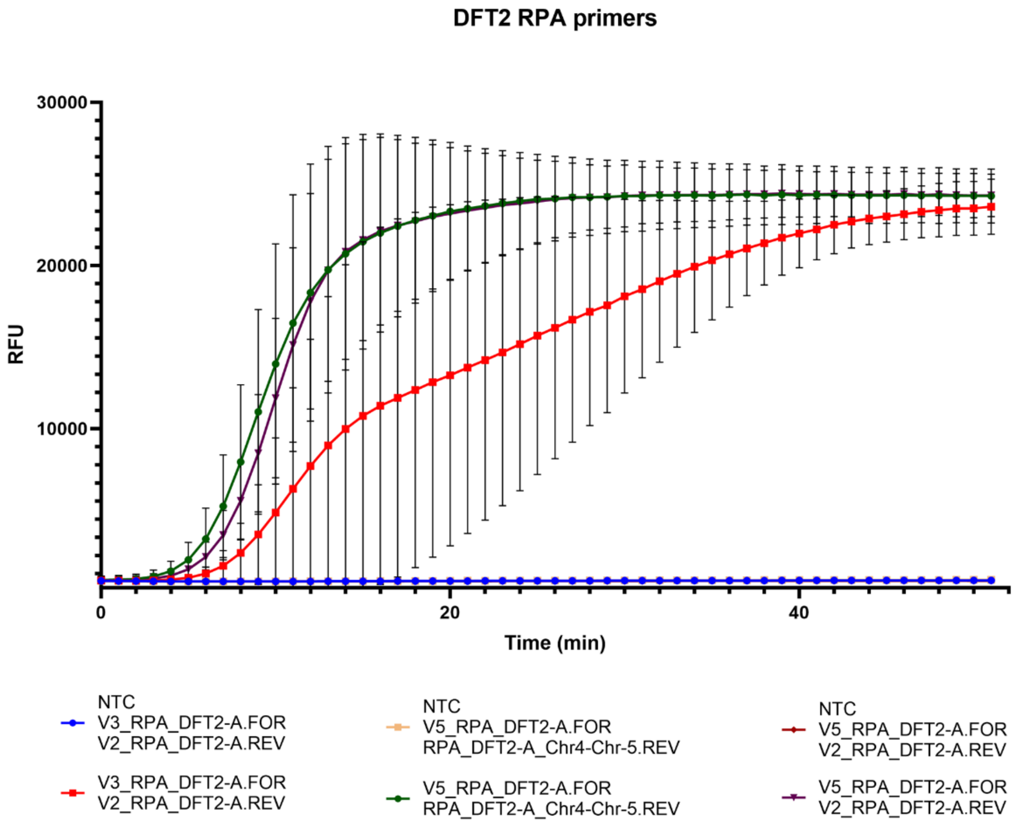


**Supplementary Figure 3. Fluorescence kinetics of top performing DFT2 RPA primers.**

Fluorescence kinetics (relative fluorescence unit, RFU) over time (minute, min). Three pairs of RPA primers for DFT2 were assessed to identify the best performing primer combination. These primers were paired with DFT2 gDNA or water as a no template control (NTC).

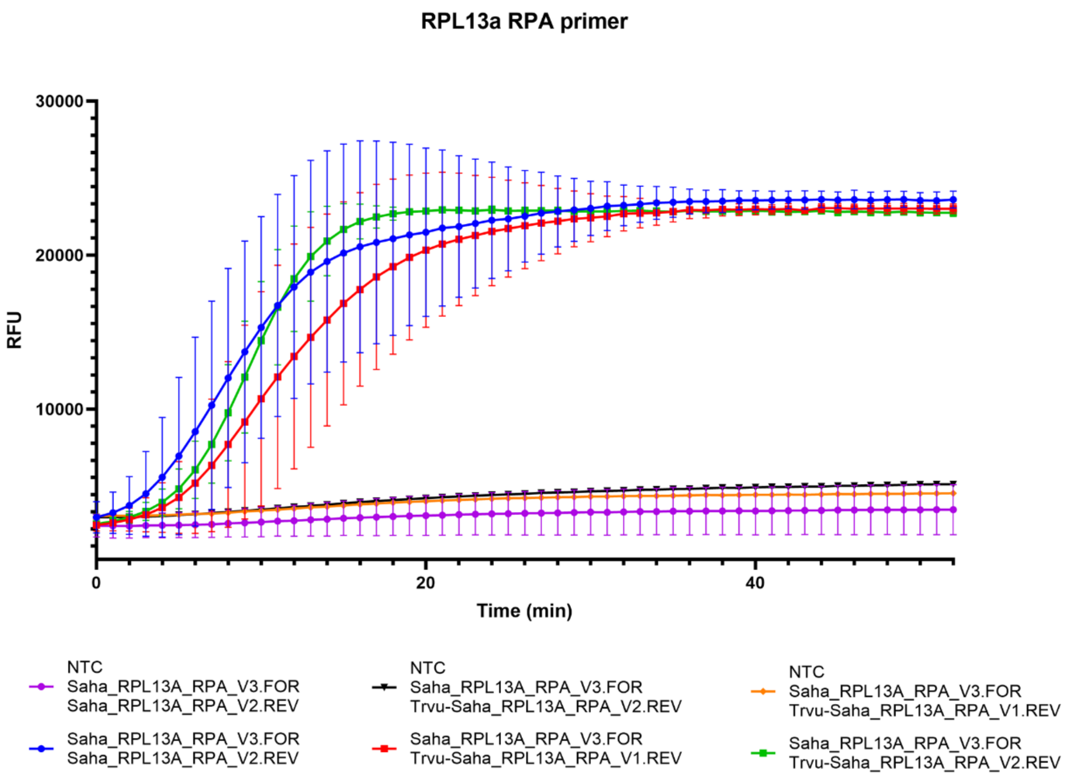


**Supplementary Figure 4. Fluorescence kinetics of top performing RPL13a RPA primers.**

Fluorescence kinetics (relative fluorescence unit, RFU) over time (minute, min). Three pairs of RPA primers for RPL13a were assessed to identify the best performing primer combination. These primers were paired with fibroblast gDNA or water as a no template control (NTC).

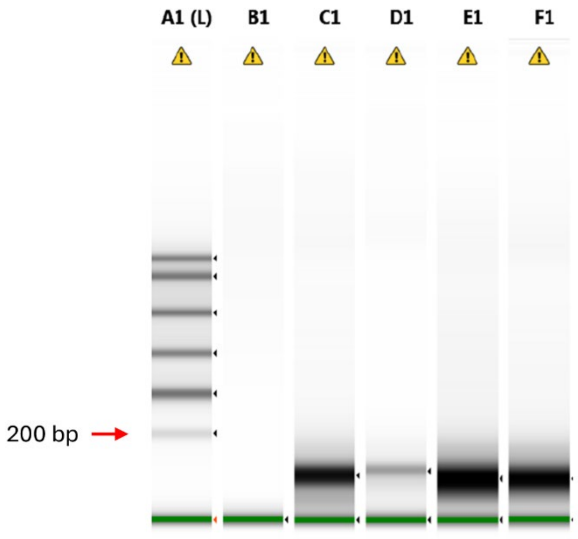


**Supplementary Figure 5.** ***In vitro*** **transcribed crRNA quality assessment.**

*In vitro* transcribed ZIKV and three RPL13a crRNA were assessed for their quality and integrity using The Agilent TapeStation system equipped with an RNA screen cassette. A1 (L), ladder; B1, no template control; C1, ZIKV crRNA; D1, RPL13a V1 crRNA; E1, RPL13a V2 crRNA; F1, RPL13a V3 crRNA. All crRNAs are 64 bp in size.

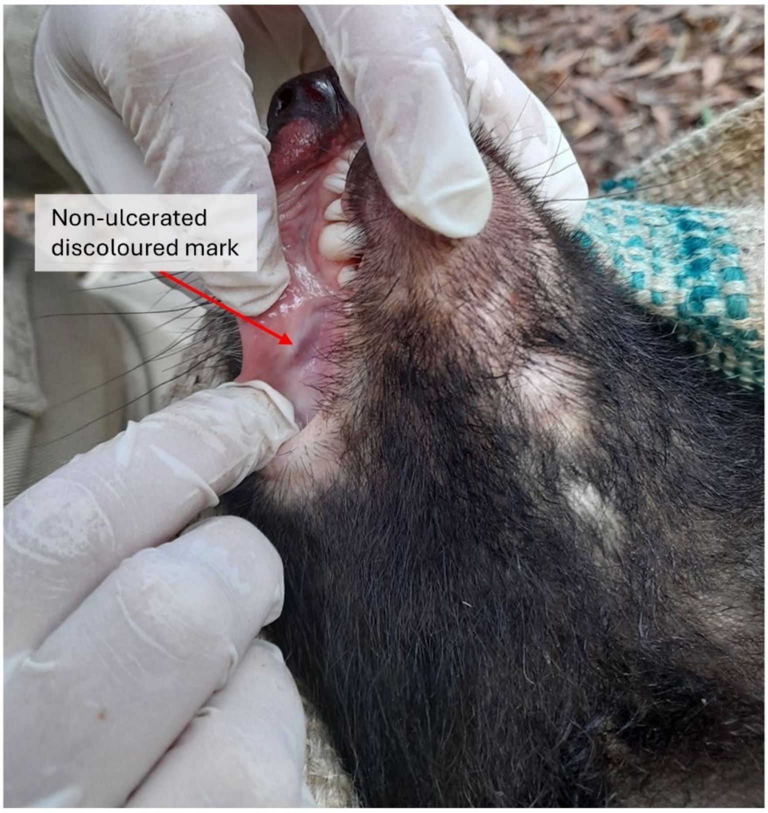


**Supplementary Figure 6. Discoloured marked observed on Fajita's mucosal tissue.**

Picture of Fajita's non-ulcerated discoloured mark as indicated by an arrow.
